# Cooperation shapes bacterial niche breadth evolution and patterns of diversification

**DOI:** 10.1101/2024.10.05.616009

**Authors:** Chunhui Hao, Naoki Konno, Makoto Ito, Laurence J. Belcher, Wataru Iwasaki, Stuart A. West

## Abstract

Bacteria exhibit varying niche breadths, with generalists thriving in diverse environments and specialists confined to specific habitats. This variability reflects the adaptability of bacteria to their environment and may influence their speciation and extinction rates. We used phylogenetic causal inference and diversification analysis techniques to investigate the influence of cooperation on bacterial niche breadth evolution and patterns of diversification across 25,785 species. We found: (1) a positive correlation between the proportion of genes for cooperation and niche breadth; (2) a decreased proportion of genes for cooperation promotes niche contraction; and (3) species with a higher proportion of genes for cooperation show increased speciation and extinction rates when their niche breadths are narrower. Our study highlights the role of genes for cooperation in shaping both niche breadth and diversification of bacteria, underscoring their critical function in maintaining the ecological versatility and diversity of bacteria.

## Introduction

Bacteria exhibit remarkable diversity and adaptability, thriving in a wide range of environments^1–5^. The concept of niche breadth, which refers to the variety of habitats a species can inhabit, can help us understand the global patterns in bacterial diversity. Certain bacterial species are “habitat generalists,” capable of living in a diverse range of environments, while others are “habitat specialists,” restricted to very specific niches^6–8^. Unravelling the mechanisms that can explain this variability in bacterial niche breadth remains a major challenge for evolutionary and ecological research^9–13^.

One factor that may play a significant role in bacterial niche breadth evolution is cooperation. Bacteria produce and release a diversity of factors that facilitate growth for the local population of cells, and hence act as cooperative ‘public goods’^14–17^ (Figure 1a). These factors play a key role in the ability of bacteria to grow in different environments. For instance, iron scavenging siderophore molecules can increase iron uptake in environments where this mineral is scarce^18^. Previous studies proposed that cooperation facilitates niche expansion and found evidence to support this hypothesis, showing that species inhabiting more habitats possess more genes for extracellular proteins^19,20^. These proteins are likely to behave as cooperative ‘public goods’ because they will often diffuse away from the producing cell and so their benefit is shared with neighbouring cells^21–24^.

**Figure 1.**
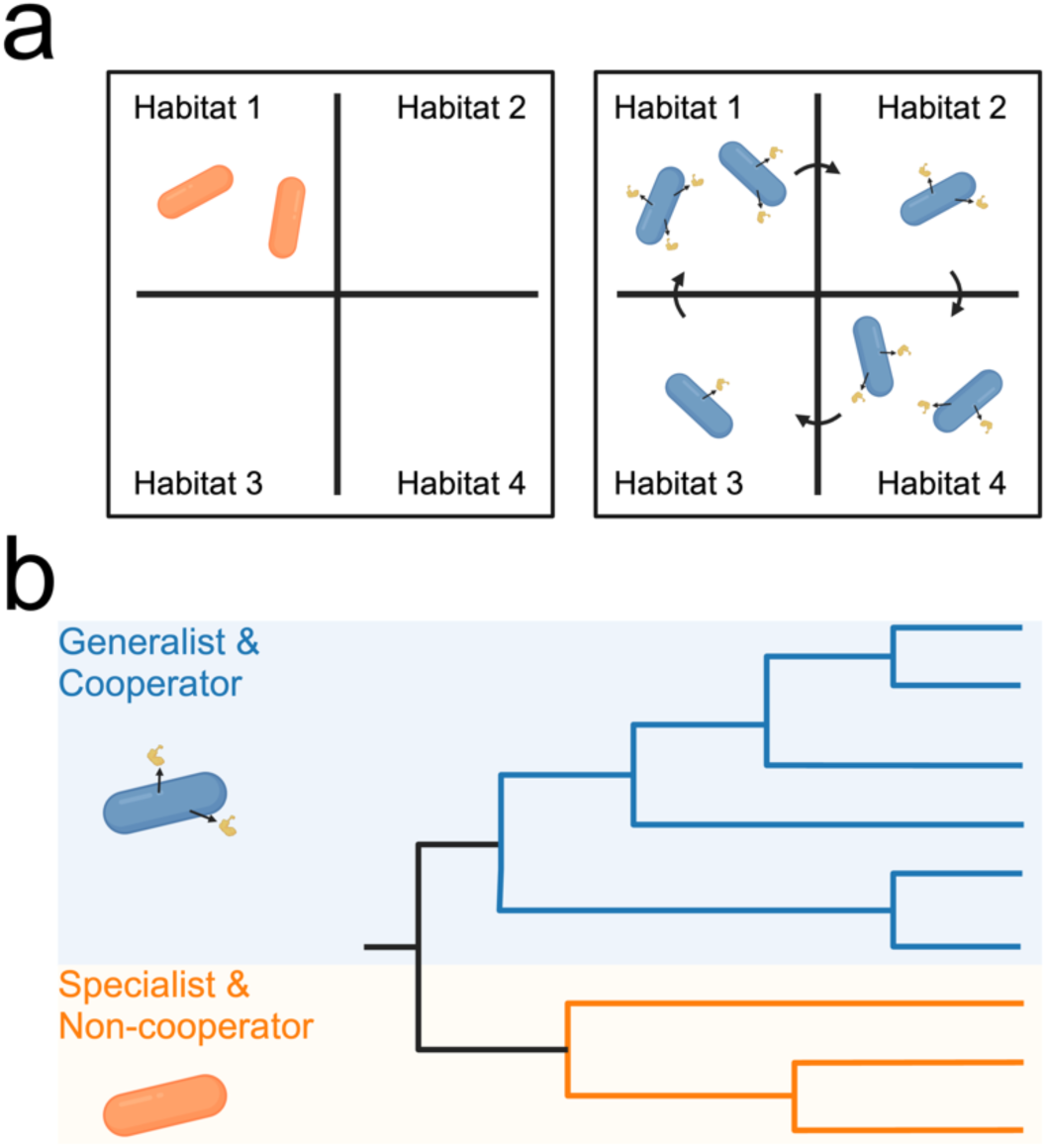
(a) Cooperative bacteria might have broader niche breadths because the production of cooperative public goods (blue cells secreting golden molecules) facilitates adaptation to different habitats. In contrast, non-cooperative bacteria (orange cells) might only adapt to specific habitats. (b) Generalists may have higher diversification rates compared to specialists. This is because generalists are exposed to diverse environments, each imposing unique selective pressure that facilitates the emergence of new species, and are better at surviving in those environments. Consequently, the diversity (number of species within a clade of the tree) of generalists might be higher than that of specialists. However, since generalists were proposed to carry more genes for cooperative traits, it is also possible that bacterial species with more such genes might have higher diversification rates. It remains unclear whether cooperation influences bacterial diversification rates directly or indirectly through its effect on niche breadth evolution.

The hypothesis that cooperation favours niche expansion could however be incorrect. An alternative explanation for the data is that causation is in the opposite direction — an initial expansion into new niches could increase the utility of genes for cooperation, making them more prevalent in generalists. In this case, mechanisms such as metabolic flexibility or bacterial motility might actually be the main drivers for niche expansion, with increased cooperation emerging as a by-product of niche expansion^13,25–27^. Another possibility is that the relationship between bacterial niche breadth and cooperation might not be causal at all, but instead could be influenced by a third, unidentified variable correlated with both^28,29^. These alternative explanations can only be distinguished by testing for different hypotheses about the causal relationship between bacterial cooperation and niche breadth evolution.

Niche breadth evolution and cooperation could also play key roles in shaping bacterial diversification^30^. Previous studies have suggested that species with broader niche breadths may experience higher diversification rates, likely due to their exposure to diverse environments, each imposing unique selective pressure^9,31^. These studies, however, have primarily focused on niche breadth, leaving open the possibility that the observed relationship between niche breadth and diversification rates is either an artefact or mediated by cooperation (Figure 1b). For instance, a higher rate of cooperation could directly facilitate diversification, resulting in an indirect correlation between niche breadth and diversification.

We investigated (1) the role of bacterial cooperation in niche breadth evolution and (2) the influence of both niche breadth and cooperation on bacterial diversification. We used improved ways to characterize the niche breadths and identify genes for cooperative traits in 25,785 bacterial species using their genomic data. Based on these data, we applied phylogeny-based analytical approaches to examine correlations between bacterial cooperation and niche breadth evolution, and to test various causal hypotheses underlying these correlations. Finally, we explored the collective effect of niche breadth and cooperation on shaping bacterial diversification rates and elucidated the taxonomic variations in these patterns.

## Results

### Niche breadth assessment

We obtained the genomes of 62,291 bacterial species from the Genome Taxonomy Database (GTDB^32^). To assess their niche breadth, we first inferred the habitat preferences of species in our dataset possessing full-length 16S rRNA gene sequences. We successfully annotated the habitat preferences for 32,262 species using ProkAtlas, a database containing environmental 16S rRNA gene sequences from 114 distinct habitats like soil, human gut, and seawater^33^ (Figure E1).

A potential problem with such habitat classifications is that they could be influenced by human biases. Environments that we classify as distinct could be functionally similar habitats for bacteria^34^. To address this, we used a method that grouped similar habitats into “habitat clusters” based on species composition^9^. If two habitats, say A and B, are inhabited by mostly the same set of species, they were grouped together as a single habitat cluster. Conversely, if few species are shared between habitats A and B, they were treated as independent habitats. Using this approach, we reorganized the original 114 habitats into 26 habitat clusters (Figure 2; Table S3; see in Methods). The habitat clustering confirmed some intuitive groupings, such as clustering chicken gut, human gut, and pig gut into a single habitat cluster. But it also revealed some non-trivial associations. For example, the method grouped paper pulp and insect gut into a single habitat cluster, suggesting similarities between these two environments in terms of microbial adaptation. Some habitats, such as plastic and biofilter, could not be grouped with other habitats, suggesting that these habitats should be considered as independent habitats (Figure 2).

**Figure 2.**
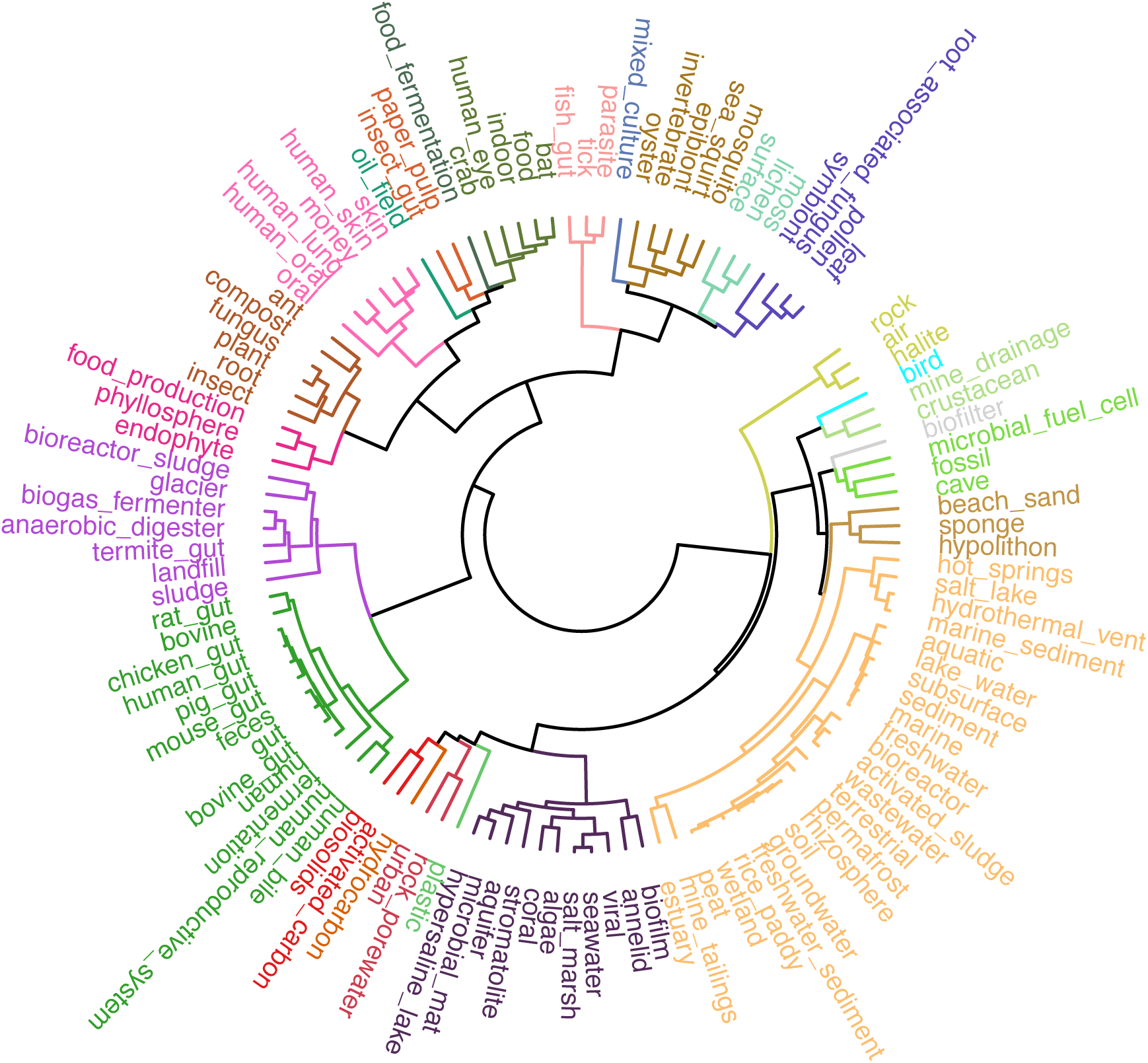
Assessment of bacterial niche breadths. Habitat preferences for each of the 32,262 species in our dataset were inferred using ProkAtlas, which contains 16S rRNA gene sequences from 114 distinct habitats. These habitats were subsequently grouped into “habitat clusters” based on similarities in species composition. Species’ habitat preference annotations were then updated accordingly. The 114 original habitats from the ProkAtlas database were clustered into 26 habitat clusters. The branches and labels of different clusters in this dendrogram were coloured differently. Niche breadth for each species was quantified by the number of habitat clusters in which they were found.

We defined the niche breadths of each species as the number of habitat clusters in which they were found. Species found in more habitat clusters are considered to have broader niche breadths (i.e., generalists) than those found in fewer habitat clusters (i.e., specialists; Figure E2a).

### Species with broader niche breadths carried a higher proportion of genes for cooperative traits

Previous studies examined the relationship between niche breadth and the carriage of genes encoding extracellular proteins^19,20^. While this approach captured genes for many cooperative behaviours, it overlooked genes which code for more complex cooperative functions, such as when a protein is intracellular but involved in producing a cooperative molecule^35^. We therefore identified genes for cooperative traits with an improved approach. First, genes were designated as “cooperative” if they were tagged with one or more of the following cooperative behaviours commonly found in bacteria: biofilm formation, quorum-sensing, secretion systems, siderophore production and usage, and antibiotic degradation^35,36^. Second, we also counted genes encoding extracellular proteins as genes for cooperative traits. Using this definition, we identified genes for cooperative traits by analysing the protein sequence data of species in our dataset using the KEGG functional annotation database^37,38^ (Figure E1). 579 KEGG-defined orthologous groups (KOs) were defined as cooperative, and genes assigned to these KOs were considered as encoding cooperative traits (Table S4).

We successfully identified genes for cooperative traits in 25,785 species of our dataset (Table S5). Given that species having broader niche breadths typically have larger genome sizes (Figure E2c), we used the proportion of genes for cooperative traits within each species’ representative genome, rather than the number of such genes, as an indicator of the level of cooperative gene carriage for that species while controlling for its genome size. The proportion of genes for cooperation ranged from around 0 to 0.07 across all species (Figure E2b).

We found a positive correlation between the proportion of genes for cooperative behaviours and niche breadth across species (MCMCglmm^39^: n = 25,785 species; pMCMC < 0.001; R^2^ of fixed effect = 0.051; Figure 3a, b; Table S1). In addition, we also found the same pattern when analysing different types of genes for cooperative traits separately (MCMCglmm: n = 25,785 species; pMCMC < 0.001 for all categories of genes for cooperation; Figure 3b, c; Table S1).

**Figure 3.**
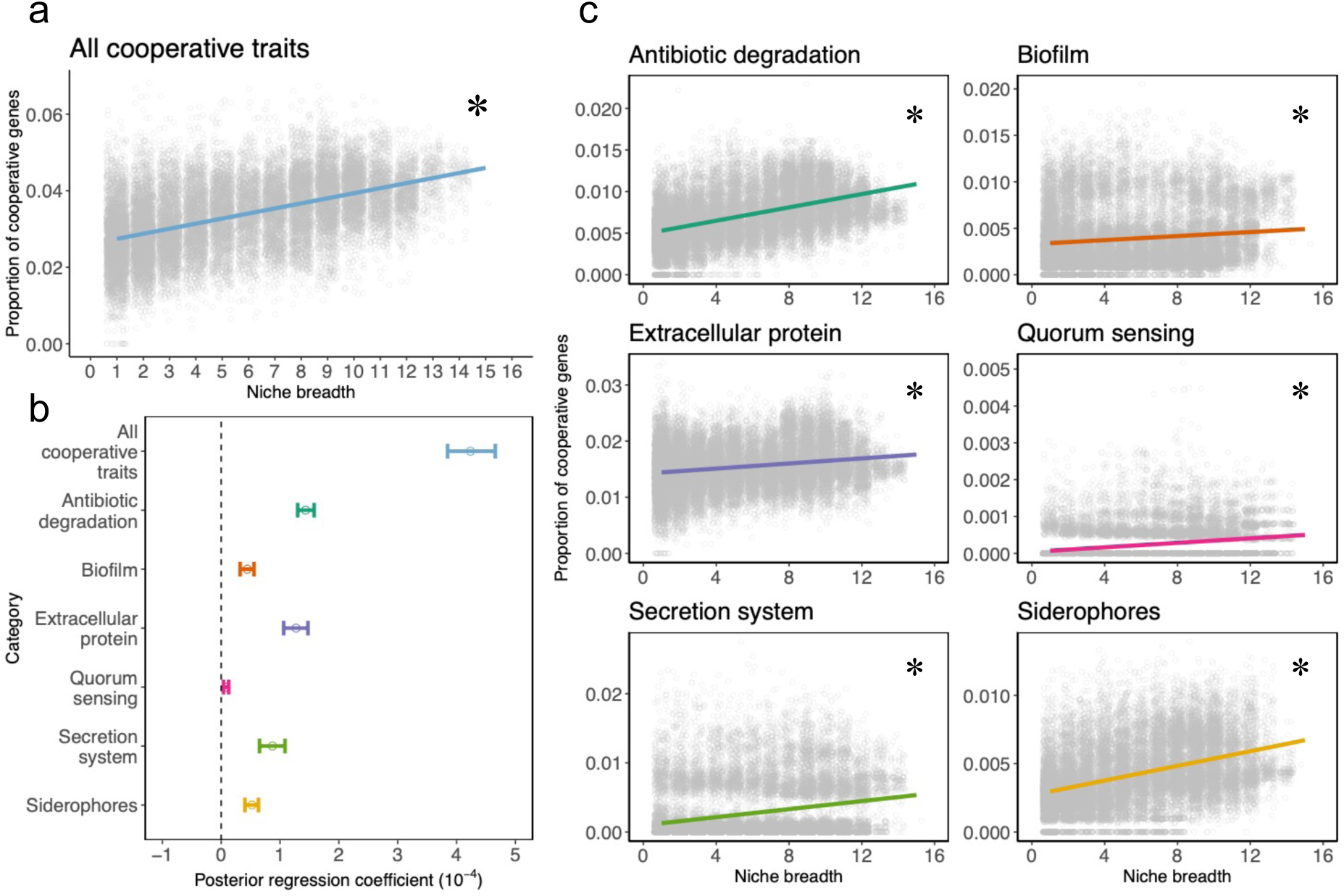
Correlation between bacterial cooperation and niche breadth. (a) Using MCMCglmm, we found a positive correlation between the proportion of genes for cooperative traits and niche breadth, defined as the number of habitat clusters a species is found in. The asterisk denotes significant correlation. (b) The mean values (dot) and 95% credible intervals (horizontal bar) of posterior regression coefficients for these correlations. (c) A consistent positive correlation between the proportion of genes for cooperation and bacterial niche breadth was observed across various functional categories of genes for cooperation. Asterisks denote significant correlations.

We conducted several tests to confirm the robustness of our findings. We found the same pattern when examining the relationship between the number of genes for cooperative traits and niche breadth (rather than proportion; Figure S1), and when using different methods (or thresholds) for clustering habitats (Figure S2 & 4). Finally, we considered the potential confounding effects of genes that may have coevolved with genes for cooperative traits, particularly those sharing the same metabolic pathway, which are prone to simultaneous gains or losses with genes for cooperative traits due to their functional interdependencies^40–42^. We found that including the proportion (or number) of coevolved genes as a covariate did not change the correlation between the proportion (or number) of genes for cooperative traits and niche breadth (Figure S8).

### Exploring the causal link between cooperation and niche breadth evolution

Our observation that habitat generalists possess a higher proportion of genes for cooperative traits is consistent with previous studies^10,20^. This correlation could reflect various causal relationships, for instance, cooperation could have facilitated niche expansion, or niche expansion could have favoured the evolution of cooperation. To test different causal hypotheses, we examined the chronological order of evolutionary events, considering earlier events lead to subsequent ones.

To carry out causal inference, we classified species as habitat generalists or specialists based on the number of habitat clusters they occupied. Specialists, by definition, have the narrowest niche breadths, so we defined species found in only one habitat cluster as specialists. To identify generalists, we compared the observed pattern to the expected distribution of niche breadths across species. We generated an expected distribution of niche breadths using a permutation approach (n = 10,000) that involved randomly assigning species to different habitat clusters while preserving the original species count in each cluster (see in Methods). Thresholds for classifying species as generalists were set at points where the observed count of species with a particular level of niche breadth significantly exceeded the expected count (Figure 4e; Table S2). Accordingly, species found in 8 or more habitat clusters were classified as generalists, while those in between were classified as having “intermediate” niche breadths (i.e. 2-7 clusters; Figure 4e). Out of the 25,785 species in our dataset, 4,230 were classified as specialists, 12,975 as intermediate species, and 8,580 as generalists (Table S5).

**Figure 4.**
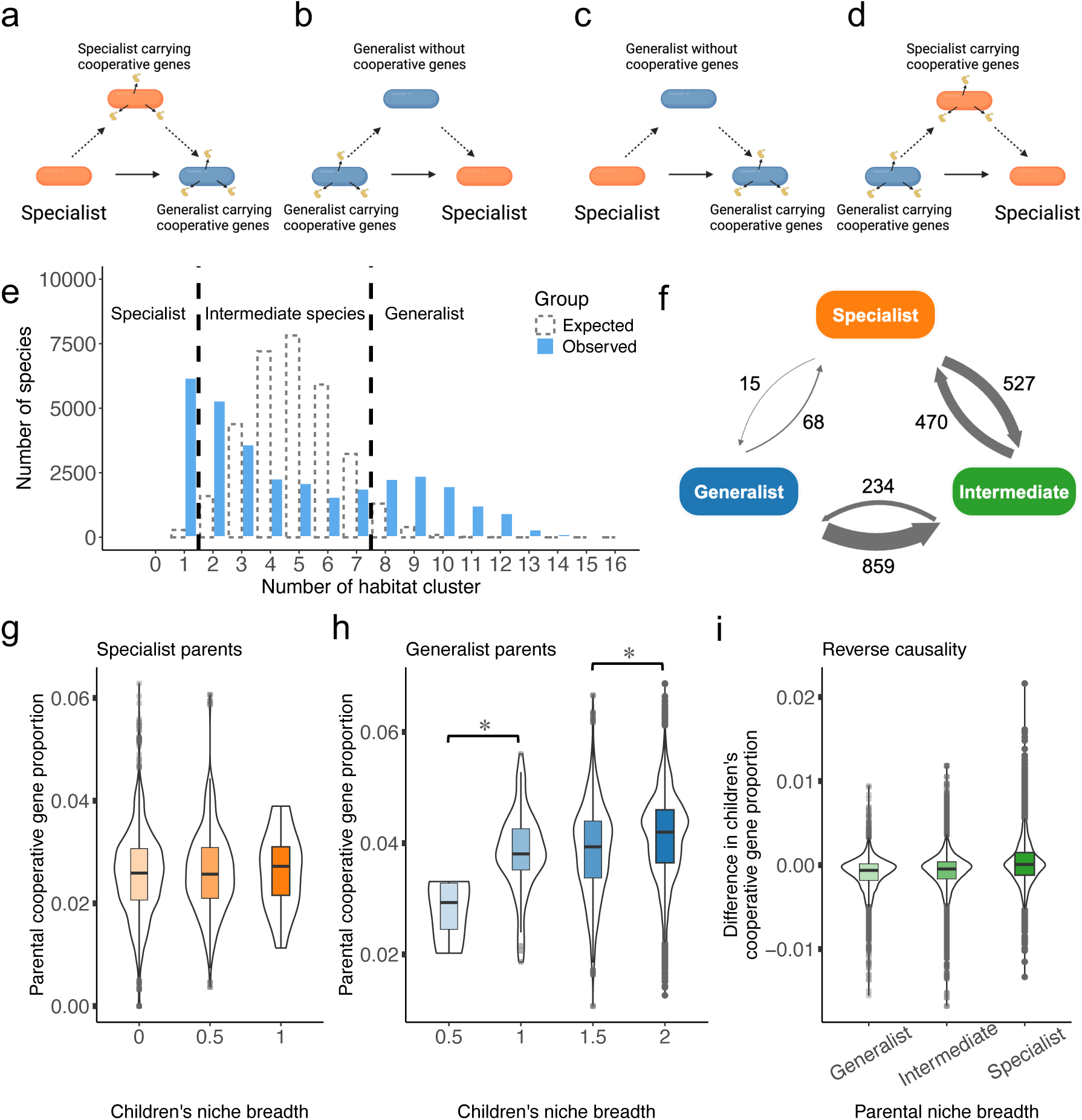
Testing causal hypotheses for the relationship between bacterial cooperation and niche breadth evolution. (a-d) Illustration of four potential causal relationships between changes in cooperative gene carriage and changes in niche breadth. Orange cells signify specialists, while blue cells represent generalists. Cells secreting golden molecules represent those carrying genes for cooperative traits. Solid arrows depict observed transition scenarios, while dashed arrows suggest the possible order of evolutionary events. For instance, panel (a) illustrates a scenario where a specialist first acquires genes for cooperation, then transits to a generalist carrying such genes. This signifies the direction of causation where having a higher proportion of genes for cooperation facilitates generalization. (e) Specialists were defined as species found in only one habitat cluster. Afterwards, the observed distribution of niche breadths was then compared against an expected distribution from 10,000 permutations to found generalist species. Those in 8 or more habitat clusters were identified as generalists, and species found in between were classified as having intermediate niche breadth. (f) Number of niche breadth transition events. Arrows indicate the direction of transition, with line widths proportional to the number of transition events. Only the transition events that occurred between each ancestral species (parent) and its two direct descendants (children) were counted to simplify the analysis. (g-i) Results of causal inference. (g) The prevalence of genes for cooperation in specialist ancestors did not significantly influence the niche expansion of their descendants, indicating that higher levels of cooperation are not a driving force for becoming generalists. (h) Generalist ancestors with descendants experiencing substantial niche contraction had lower proportions of genes for cooperation, suggesting that a reduced prevalence of genes for cooperation might facilitate specialization. (i) No significant relationship was observed between the niche breadth of ancestral species and the change in the proportion of genes for cooperation among their descendants, suggesting that transitions to either generalist or specialist do not affect the carriage of genes for cooperation in the offspring.

We formulated four potential causal scenarios based on the chronological order of events, where either the level of cooperation influences the evolution of niche breadth or the reverse (Figure 4a-d). To infer the earlier evolutionary events, we applied ancestral state reconstruction to recreate the gene content profiles and niche breadths for ancestral species (internal node), based on data from extant species (tip) and their known phylogenetic relationships^43^ (Figure E3). We reconstructed the gene profiles and niche breadths for 24,912 ancestral nodes, which included 3,994 specialists, 11,683 species with intermediate niche breadth, and 9,235 generalists. The ancestral niche breadths for 872 nodes were undetermined (Figure E4a, Table S6). We only compared the evolutionary transitions between each ancestor (parent) and its direct descendants (children) to maintain analytical clarity (Figure E3). Each parent had two children, which could be specialists (quantified as 0), intermediate species (quantified as 0.5), or generalists (quantified as 1). For each parent, we summed niche breadths for both children yielded possible children’s niche breadth scores of 0 (two specialists), 0.5 (one specialist, one intermediate species), 1 (two intermediate species, or one specialist one generalist), 1.5 (one generalist, one intermediate species), or 2 (two generalists).

We found support for the causal hypothesis that a lower proportion of genes for cooperative traits facilitated niche contraction (Figure 4b). We classified 927 transition events from generalists to species with narrower niche breadths, comprising 68 events from generalists to specialists, and 859 events from generalists to intermediate species (Figure 4f). We found that generalist parents whose children underwent significant niche contraction had a lower proportion of genes for cooperation (MCMCglmm: n = 9,173 generalist ancestral species; 1 vs. 0.5: pMCMC < 0.001; 1.5 vs 1: pMCMC = 0.663; 2 vs 1.5: pMCMC < 0.001; Figure 4h, Table S1).

In contrast, we found no support for the hypothesis that a higher proportion of genes for cooperation promote niche expansion (Figure 4a). The total number of transition events from specialists to species with broader niche breadths was 542, comprising 527 events from specialists to intermediate species, and 15 events from specialists to generalists (Figure 4f). We found that the proportion of genes for cooperation in specialist parents did not significantly affect whether their children expanded their niches (MCMCglmm: n = 3,832 specialist ancestral species; pMCMC = 0.637; Figure 4g, Table S1).

We also did not find support for the hypotheses that niche breadth influenced the level of cooperation (Figure 4c, d). We calculated the change in the proportion of genes for cooperative traits between each ancestral species and the average for its two children. Our analyses showed no significant correlation between the ancestral species’ niche breadth and the change in the proportion of genes for cooperative traits in their descendants (MCMCglmm: n = 24,590 ancestral species; pMCMC = 0.796; Figure 4i, Table S1). This suggested that neither becoming a generalist nor becoming a specialist influenced the evolution of the proportion of genes for cooperation in the offspring.

To test the robustness of our results we also carried out a transition rate analyses, which allows us to examine whether the evolution of two traits is correlated, as well as the underlying direction. We again found support for the hypothesis that a lower proportion of genes for cooperation facilitated specialization (Figure 4b). First, the transition rate from broader to narrower niche breadths (specialization) was higher than the transition rate from narrower to broader niche breadths (generalization). Second, the transition rate from broader to narrower niche breadths was higher in species with fewer genes for cooperation compared to those with more such genes (Figure S9).

### Genes for cooperation undergo more frequent intraspecific gains and losses

Why would a lower proportion of genes for cooperative traits favour niche contraction? We hypothesized that genes for cooperative behaviours act as a maintenance mechanism for relative generalists, that allows them to maintain broad niches. Losing these genes undermines this ability, pushing the species towards relative specialists. In contrast, for specialists, while the potential benefits of genes for cooperative behaviours, such as enhanced resource utilization, might theoretically facilitate niche expansion, the associated costs, such as the energy expenditure for synthesizing these genes for cooperative behaviours, or the vulnerability to exploitation by ‘cheaters’, also act as selective pressures for their deletion^15,21,44,45^. This would lead to a rapid turnover in the acquisition and loss of genes for cooperative traits over relatively shorter-term, such that they do not significantly impact the longer-term evolutionary transition from being a specialist to a generalist.

We tested this hypothesis by comparing the rate at which genes for cooperation are gained and lost both across species (interspecific) and within species (intraspecific). The intraspecific rate represents gain and loss over relatively shorter evolutionary timescales, whereas the interspecific rate represents these processes over longer timescales^46^. We would expect that genes for cooperation undergo frequent gains and losses within species, but not across species.

To estimate the rate at which genes for cooperation are gained and lost within a species (intraspecific), we examined bacterial pangenomes. A pangenome comprises all the genes found across different strains of a species^47^. Within this, some genes are ubiquitous across all strains (core genes), and some are only present in a subset of strains (accessory genes), making them more prone to gains and losses compared to core genes^40^ (Figure 5a). If genes for cooperation are more likely to undergo frequent gains and losses within species, then we would expect them to be more likely to be in the accessory genomes.

**Figure 5.**
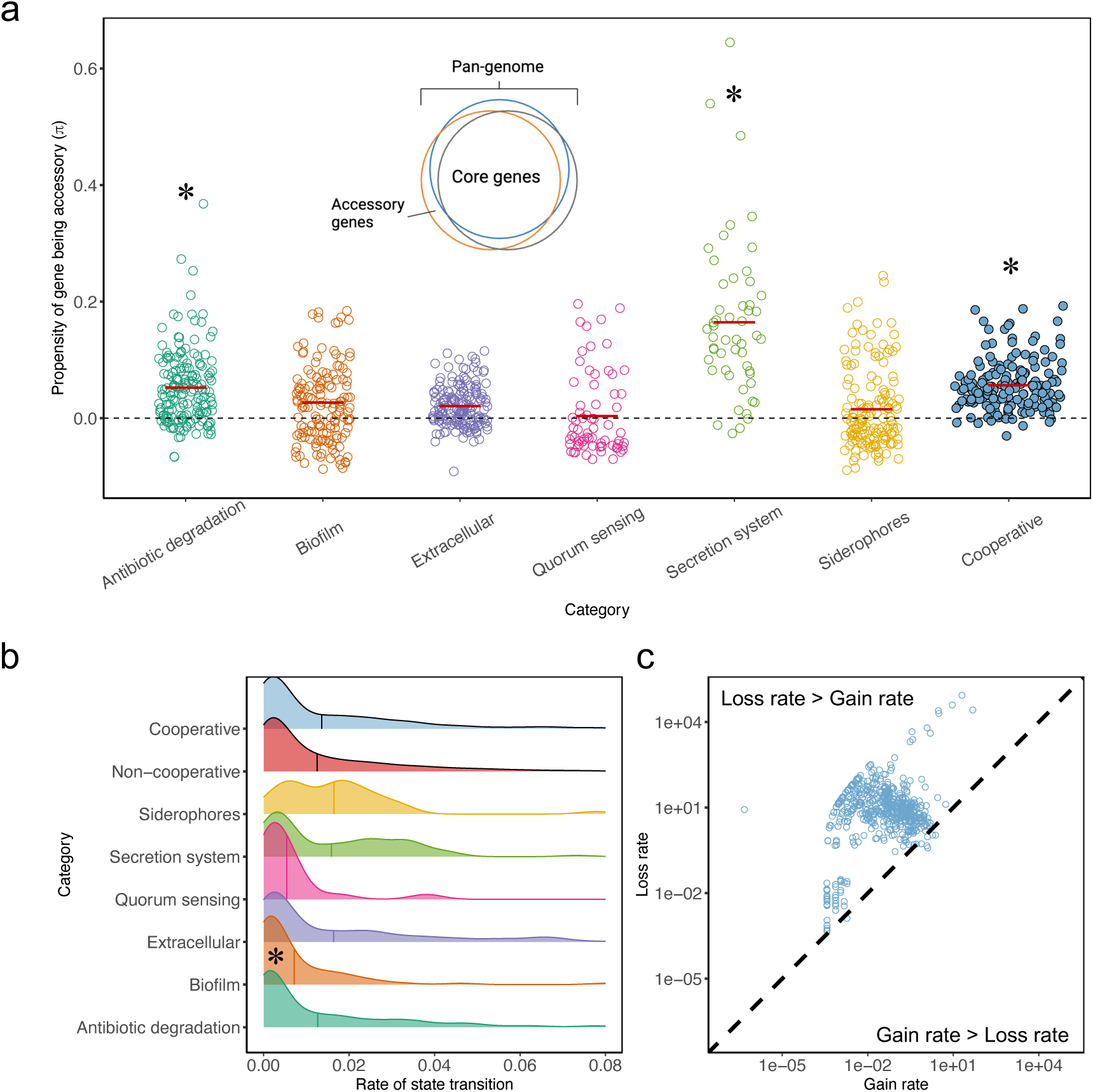
Gain and loss dynamics of genes for cooperative traits. (a) At the intraspecific level, the gain and loss rates of genes for cooperation are indicated by the propensity of a given gene for cooperation to be an accessory gene in a pangenome, meaning these genes are present only in a subset of strains of a species. The metric π is used to quantify the propensity of a gene being an accessory gene in a pangenome compared to random expectation. Each dot represents π for a specific gene type within a species, with red lines indicating the average π for each gene type across all species. Genes for cooperation were significantly more likely to be accessory genes, suggesting a higher intraspecific gain and loss rate for these genes, especially pronounced in genes related to secretion systems and antibiotic degradation. (b) The gene gain and loss rate across species (interspecific) is measured by the rate of state transitions for each KO. Ridgeline plots visualize the interspecific gene gain and loss rates, displaying the distribution of these rates for each gene type. The central line in each plot indicates the mean state transition rate for each gene type. No significant differences were observed between cooperative and non-cooperative KOs. However, biofilm formation-related KOs displayed significantly lower transition rates compared to non-cooperative KOs. (c) The gain rates are plotted against the loss rates for each cooperative KO on a log-log scale. Each dot represents the rate of a cooperative KO. The dashed line indicates where gain and loss rates are equal. The upper-left region shows where loss rates exceed gain rates, while the bottom-right region shows where gain rates exceed loss rates. For the majority of cooperative KOs, loss rates were found to be larger than gain rates.

We reconstructed pangenomes for 171 bacterial species for which at least 100 high-quality genomes were available in GTDB (Figure E6, Table S9). To quantify the susceptibility of genes for cooperation to gains and losses, we introduced an index π, which measures the propensity of a gene for cooperation being an accessory gene in a pangenome relative to what would be expected by chance. The higher the value of π for genes for cooperation, the more they tend to be accessory genes within bacterial pangenomes (see in Methods). In support of our hypothesis, we found that genes for cooperative traits were significantly more likely to be accessory genes (MCMCglmm: n = 171 species; pMCMC < 0.001; Figure 5a; Figure E6; Table S1). This trend was notably stronger for genes associated with secretion system and antibiotic degradation (MCMCglmm; secretion systems: pMCMC = 0.037; antibiotic degradation: pMCMC = 0.0132; Figure 5a; Table S1), but not for other functional categories of cooperative genes.

To measure gene gain and loss rate across species, we estimated the rate of state transitions for each KO per phylogenetic branch by conducting ancestral state reconstruction. Specifically, gene gains were denoted by transitions from absence to presence, while gene losses were denoted by transitions from presence to absence (Figure E5; Table S7). We found no significant difference in the state transition rates between cooperative versus non-cooperative KOs (permutated Kruskal-Wallis test; chi-squared = 3.5859, p-value = 0.060; Figure 5b).

The pattern did however vary with different types of cooperative trait. Biofilm formation-related KOs had significantly lower rates of state transitions compared to non-cooperative KOs, and those related to the formation of extracellular proteins and siderophores (Dunn’s Test: p-values were adjusted by Bonferroni correction; biofilm vs. private: p-value = 0.015; biofilm vs. extracellular: p-value = 0.0004; biofilm vs. siderophore: p-value = 0.0003; Figure 5b; Table S2). In addition, we found that cooperative KOs were more prone to gene loss than to gene gain, as demonstrated by their significantly higher rates of gene loss (permutated Kruskal-Wallis test; chi-squared = 614.39, p-value < 0.001; Figure 5c; Table S8).

Overall, our results suggested that genes for cooperative traits undergo more frequent gains and losses on a shorter time scale (within species), but not on a longer time scale (across species). These findings are consistent with our hypothesis that the cost of genes for cooperative traits leads to rapid turnover in their acquisition and loss, so that they have a limited impact on the longer-term evolutionary transition from specialists to generalists.

### Cooperation shapes speciation and extinction rates

Finally, we investigated the impact of niche breadth and cooperation on bacterial diversification rates. We fitted rate parameters of character-dependent speciation (λ) and extinction (μ) models (SSE models) using the data of two traits (cooperation & niche breadth) and the phylogenetic tree. This approach involved comparing the fit of a null model, where the trait evolves under a continuous-time Markov *k*-state (Mk) model and the tree grows under a constant birth-death process uninfluenced by the traits, to a model where speciation and extinction rates depend on the values of the traits^48^.

Fitting character-dependent SSE models required binarizing both the proportion of genes for cooperative traits and niche breadth. We classified species with values below the median proportion as having “lower” levels of cooperation and those above as having “higher” levels of cooperation. For niche breadth, species found in fewer than 6 habitat clusters were categorized as having “narrower” niche breadth, while those in 6 or more habitat clusters as having “broader” niche breadth (Figure E8). To investigate the collective effect of cooperation and niche breadth on bacterial diversification, we merged these two binary traits into a single trait with four states and applied the multi-state speciation and extinction (MuSSE) model^49^. The four states defined are: narrower niche breadth and lower cooperation (1), narrower niche breadth and higher cooperation (2), broader niche breadth and lower cooperation (3), and broader niche breadth and higher cooperation (4).

We found that fitting character-dependent SSE models using a phylogenetic tree with too many species can lead to implausible results, such as near-zero extinction rates (Figure E7). To address this issue, we fitted the MuSSE model separately for each family containing at least 100 species (Table S10). This strategy improved the accuracy of model fittings by reducing tree sizes, and enabled the comparison of diversification patterns across families while controlling for phylogeny, to eliminate the problem of pseudo-replication within lineages^50^.

Overall, the net diversification rates (speciation – extinction) at the family level did not vary significantly across different states (Figure S12, Table S1). When examining separately, we observed that species with narrower niche breadth and lower cooperation (state 1) had significantly lower speciation rates compared to those with narrower niche breadth and higher cooperation (state 2). However, there was no significant difference in speciation rates between species with broader niche breadth and lower cooperation (state 3) and those with broader niche breadth and higher cooperation (state 4). This indicated that lower cooperation levels reduce speciation rates only in species with narrower niche breadths (MCMCglmm; n = 34 families; λ_1_ vs. λ_2_: pMCMC = 0.032; λ_3_ vs. λ_4_: pMCMC = 0.357; Figure 6a; Figure E9; Table S1). Additionally, no significant differences were detected between the speciation rates of states 1 and 3 or between states 2 and 4, suggesting that niche breadth does not independently influence speciation rates (MCMCglmm: n = 34 families; λ_1_ vs. λ_3_: pMCMC = 0.221; λ_2_ vs. λ_4_: pMCMC = 0.982; Figure 6b; Figure E9; Table S1).

**Figure 6.**
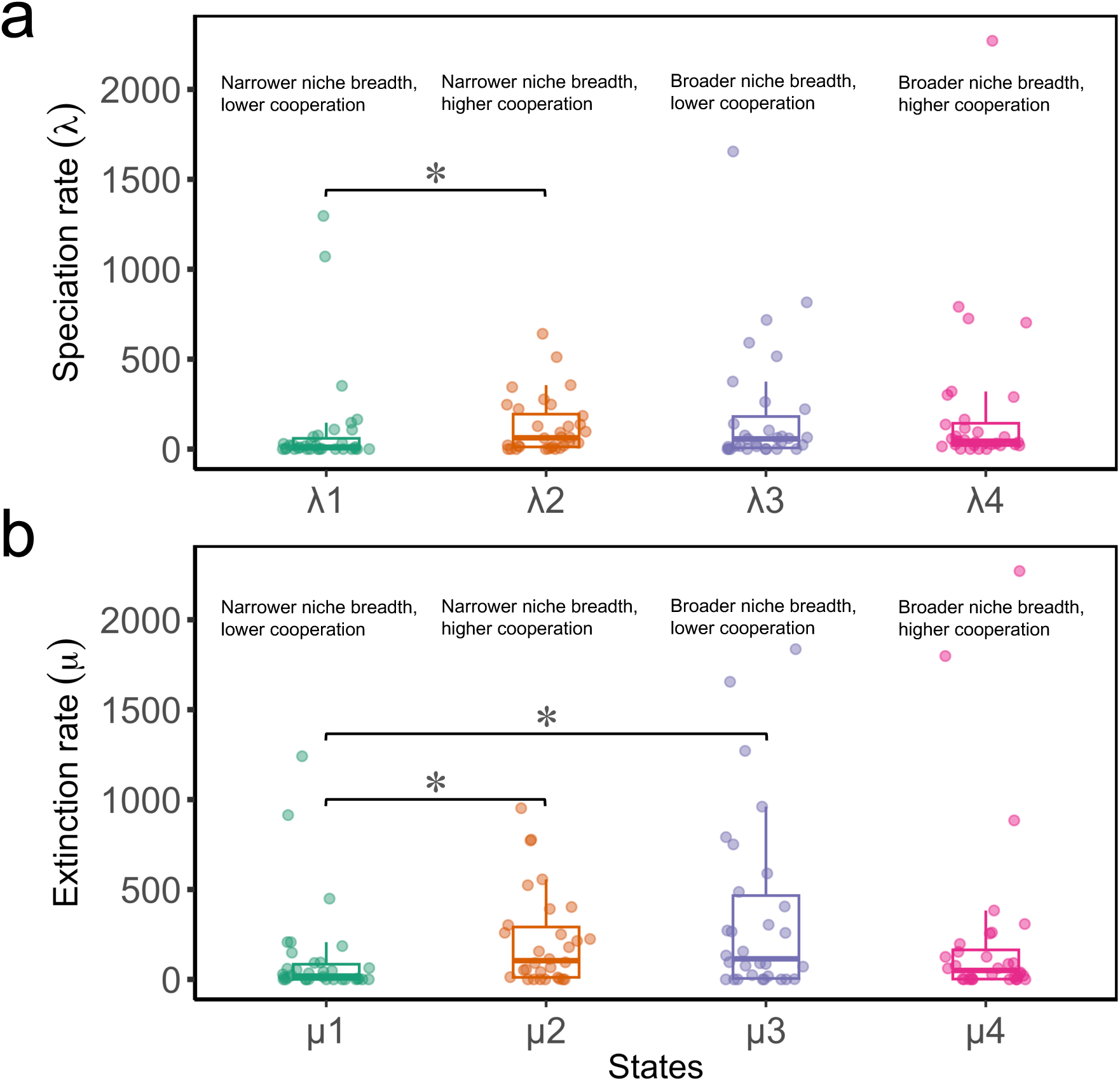
Family-wise bacterial diversification analyses. Both the proportion of genes for cooperation and the niche breadth were binarized to meet the requirement of fitting character-dependent SSE models. To assess if diversification patterns of one variable might be influenced by another, we combined the two binary variables into one with four states and explored the role of this new variable in bacterial diversification. (a) Regarding speciation rates, the rate for state 1 was significantly lower than that for state 2, indicating that species with lower cooperation levels have reduced speciation rates compared to those with higher cooperation levels when they also have narrower niche breadths. Each dot represents the rate parameter of a specific state of a trait for a given family. (b) For extinction rates, the rate for state 1 was significantly lower than that for state 2, indicating that species with lower cooperation levels have lower extinction rates compared to those with higher cooperation levels when they also have narrower niche breadths. Additionally, the extinction rate for state 1 was significantly lower than that for state 3, indicating that species with narrower niche breadths have reduced extinction rates compared to those with broader niche breadths when they also carry fewer genes for cooperative traits.

For extinction rates, we found that species with narrower niche breadth and lower cooperation (state 1) had significantly lower extinction rates compared to those with narrower niche breadth and higher cooperation (state 2). However, no significant differences were observed between the extinction rates of species with broader niche breadth and lower cooperation (state 3) and those with broader niche breadth and higher cooperation (state 4) (MCMCglmm: n = 34 families; μ_1_ vs. μ_2_: pMCMC = 0.038; μ_3_ vs. μ_4_: pMCMC = 0.644; Figure 6b; Figure E9; Table S1). This result suggested that species with lower cooperation levels experience lower extinction rates primarily when they have narrower niche breadths. Furthermore, the extinction rate for state 1 was significantly lower than that for state 3, while no significant difference was observed between the extinction rates of states 2 and 4 (MCMCglmm; n = 34 families; μ_1_ vs. μ_3_: pMCMC = 0.047; μ_2_vs. μ_4_: pMCMC = 0.636; Figure 6b; Figure E9; Table S1). This indicated a more pronounced effect of niche breadth on increasing extinction rates among species with lower levels of cooperation.

We also employed the binary-state speciation (λ) and extinction (μ) model (BiSSE) to examine the independent effects of cooperation and niche breadth on bacterial diversification^48^. When examining independently, we found that species with higher levels of cooperation tend to have higher extinction rates, while species with broader niche breadths tend to have higher speciation rates (Figure E8, Table S1). In comparison, the MuSSE models, which analyse the combined effect of cooperation and niche breadth, revealed interactions between these traits on diversification that were not captured when examining each trait independently using BiSSE models.

## Discussion

We examined the role of cooperation in bacterial niche breadth evolution and diversification. Our findings revealed that habitat generalists possess a higher proportion of genes for cooperative traits (Figure 3). Using phylogenetic causal inference techniques, we demonstrated that this positive correlation between cooperation and niche breadth is due to a causal relationship where a decreased proportion of genes for cooperative traits promotes niche contraction. This suggested that genes for cooperative traits primarily serve to maintain the niche breadth of generalists (Figure 4).

Our findings suggested that causality is in the opposite direction to the previous assumption that cooperation facilitates niche expansion, via mechanisms such as bacteria using cooperative secretions to modify the environment and extend the host range^14,19^. To understand the revealed direction of causality, we proposed that the cost of genes for cooperative traits makes them more likely to be deleted, leading to frequent acquisition and loss within species, preventing a significant impact on the long-term evolutionary transition from specialists to generalists. Our examination on the dynamics of genes for cooperation supported this explanation, showing that these genes are more likely to be accessory in pangenomes. As a result, they may undergo more frequent gains and losses within species but not across species (Figure 5).

More generally, cooperation has been found to influence the environmental choices of organisms across various taxa. In birds, cooperative breeding has been shown to facilitate the colonization of harsh environments^29^. In insects, the evolution of sociality in burying beetles has allowed them to adapt to broader elevational and temperature gradients^51^. For marine life, a study has found that eusocial sponge-dwelling shrimp occupy more sponges and have broader host ranges than their non-social sister species^52^. The major advancement made by our study, which goes beyond the current understanding of the role of cooperation, is the examination of the genetic mechanisms underlying cooperation in niche breadth evolution, and the elucidation of how a decrease in cooperation, rather than an increase, influences niche breadth evolution.

Finally, we assessed the combined effect of cooperation and niche breadth on bacterial diversification at the family level. Contrary to previous studies that found species with broader niche breadth have higher speciation rates^9,31^, our results indicated that niche breadth alone does not directly shape speciation rates. Instead, we found that cooperation increases speciation and extinction rates in species with narrower niche breadths, suggesting that it is the interaction between bacterial cooperation and niche breadth that shapes these rates (Figure 6). Although the overall diversification rates (speciation – extinction) at the family level were found not to be affected by these two traits, this is potentially because these rates were not directly fitted and were therefore prone to biases inherent in the maximum likelihood estimation process^53^. Previous research has similarly found that cooperation with symbiotic microbes has influenced diversification across insects by shaping the feeding niche of these insects^54^. Our findings, together with those from previous research, underscore the broader significance of cooperation in driving diversification, revealing its pivotal role in shaping evolutionary processes across various taxa.

## Methods

### Species selection and genome collection

We obtained a list of 62,291 species and their corresponding genomes from the Genome Taxonomy Database (GTDB^32^; release 207) on 1^st^ October 2022. At the first step, we selected species with representative genomes in GTDB that enabled the identification of full-length 16S rRNA gene sequences, yielding a list of 38,474 species. To ensure high-quality of genomes, we applied a stricter criterion than a previously proposed Minimum Information about a Metagenome-Assembled Genome (MIMAG) criterion^55^, considering genomes with completeness ≥ 95% and contamination ≤ 5% as high-quality genomes. We then used *ncbi-genome-download* scripts (version 0.3.1) to download the nucleotide and amino acid sequences for all available genomes from either GenBank^56^ or RefSeq^57^ database using the assembly entries specified in GTDB. This resulted in a dataset of 237,354 genomes with corresponding sequences for subsequent analysis.

### Habitat preferences annotation

The annotation of habitat preferences followed an approach modified from a previous study^9^. The procedure comprised three steps: (a) inferring the preliminary habitat preferences for each species using their 16S rRNA gene sequences; (b) clustering the preliminary habitats based on species compositions; (c) re-annotating the habitat preferences of each species by replacing the preliminary habitats with the newly defined habitat clusters.

#### a) Preliminary habitat estimation

To estimate the preliminary habitat preferences for each species, we first retrieved the full-length 16S rRNA gene sequences for their representative genomes from GTDB (https://data.gtdb.ecogenomic.org/releases/release207/207.0/genomic_files_reps/bac120_ssu_reps_r207.tar.gz). We then used BLAST to search these representative full-length 16S rRNA sequences against the 16S rRNA sequences in metagenome datasets containing various environmental information through the ProkAtlas online platform^33^. The sequence similarity threshold was set to 97%, and the sequence coverage threshold was set to 150 bp for conducting the ProkAtlas annotation. A species was considered present in an environment if its habitat preference score, reported by ProkAtlas, was greater than zero. We successfully estimated habitat preferences for 32,262 species out of 38,474 species from GTDB (Table S11).

#### b) Clustering habitats based on species compositions

To avoid potential bias arising from human intuition in the preliminary habitat definitions, we adopted a more objective biological approach by clustering habitats based on their species assemblages^9^. This method aimed to identify situations where two habitats, though differently defined by humans, might host the same bacterial species. For instance, if two habitats (A and B) have nearly identical sets of species, they would be clustered together. Conversely, if habitats A and B share almost no species, they would be considered independent habitats.

We initially collected a comprehensive dataset of environmental 16S rRNA sequences from the ProkAtlas database^33^. This dataset comprises 361,474 sequence fragments derived from 114 distinct habitats. To determine the species-level habitat preferences of these 16S rRNA fragments, we then mapped the fragmental sequences to full-length 16S rRNA sequences available in the SILVA database^58^ (https://www.arb-silva.de/fileadmin/silva_databases/release_138.1/Exports/SILVA_138.1_SSURef_NR99_tax_silva.fasta.gz). For the mapping process, we utilized VSEARCH^59^ (v2.22.1_linux_x86_64) to perform an all-against-all BLAST search between the ProkAtlas and SILVA sequences, setting the sequence identity threshold at 98%. As a result, we successfully matched 248,295 out of 361,474 ProkAtlas fragmental sequences to 71,574 full-length 16S rRNA sequences present in the SILVA database. Each of these full-length sequences corresponded to a unique bacterial species. This enabled us to create presence/absence profiles of bacterial species associated with every ProkAtlas habitat.

To evaluate the similarities between ProkAtlas habitats and prepare for subsequent clustering steps, we calculated environmental similarity scores based on a previous study^9^. Specifically, for any pair of habitats *H1* and *H2*, and for a sequence identity threshold *T* (ranging from 70% to 98%), the similarity score between *H1* and *H2* was calculated as the geometric mean of two proportions: (i) The proportion of full-length 16S rRNA sequences (species) in *H1* that were similar to any sequence (species) in *H2* with a sequence identity above the threshold *T*; (ii) The proportion of full-length 16S rRNA sequences in *H2* that were similar to any sequence in *H1* with a sequence identity above the threshold *T*. The sequence matching was performed with BLASTn (version 2.13.0) in an all-against-all mode between each pair of sequence sets at the threshold *T*. We allowed *T* to vary because species do not necessarily need to be identical (*T* = 98%) to exhibit similar habitat preferences. Bacterial species from higher taxonomic ranks might possess adaptive traits that enable them to thrive in the same environments, even with some genetic divergence^60^. As a result of this process, we obtained matrices of similarity scores between each pair of ProkAtlas habitats, each calculated at different sequence identity thresholds.

Finally, we conducted hierarchical clustering to group ProkAtlas habitats using the dissimilarity score (1-similarity score) at different thresholds *T*. To objectively determine the optimal clustering methods (between “ward.D”, “ward.D2”, “single”, “complete”, “UPGMA”, “WPGMA”, “WPGMC”, or “UPGMC”), the numbers of clusters (range from 2 to 113), and the sequence identity thresholds (range from 70% to 98%), we calculated the Silhouette index for various clustering results. The Silhouette index ranges from −1 to +1, with a higher value indicating better clustering performance, where objects are well-matched to their own cluster and poorly matched to neighbouring clusters. We found the optimal Silhouette index was achieved when we using “ward.D2” method with a threshold *T* = 77% and 26 habitat clusters (Figure E10, Table S13). The dendrogram of this optimal clustering can be checked in Figure 2. The analyses here was performed in R (version 4.2.1), using package *NbClust* (version 3.0.1)^61^ and package *Factoextra* (version 1.0.7)^62^.

#### c) Re-annotating the habitat preferences

Using the 26 newly generated habitat clusters, we replaced the previous 114 habitats, allowing us to re-annotate habitat preferences for species in our dataset. This re-annotation provided a more refined representation of species’ habitat preferences, based on a relatively objective clustering of similar environments. We successfully annotated 32,262 species from our initial species list with 25 habitat clusters (Table S12).

### Quantifying niche breadth

To quantify the niche breadths of every species in our dataset, we calculated the number of habitat clusters in which they were found. Species found in more habitat clusters are considered to have broader niche breadths than those found in fewer habitat clusters (Figure E2a).

To implement the causal inference, we classified the species as habitat generalists or specialists based on the number of habitat clusters they occupied. According to the definition of specialists, species with numbers of clusters equal to 1 were defined as habitat specialists. To identify habitat generalists, we compared the observed distribution of number of clusters across species to an expected distribution generated through permutations. Based on the observed species– habitat cluster association matrix, we conducted 10,000 random permutations of the species– habitat cluster associations, while preserving the species count in each cluster, to establish the expected distribution of the number of habitat clusters across species. This approach ensures that the null pattern reflects the assumption that species can potentially adapt to any habitat cluster, but each habitat has a limitation on the number of species it can support. Cut-offs for defining generalists were set where the observed species count at a particular number of clusters significantly exceeded the expected count. As a result, species found in 8 or more habitat clusters were classified as generalists. In addition, species with niche breadths between specialists and generalists were defined as intermediate species (Figure 4e).

### Identifying genes for cooperative traits

We adapted a method from a previous study^36^ to define genes for cooperative traits based on functional annotations. Specifically, genes were considered “cooperative” if they were annotated with at least one of the five well-known forms of bacterial cooperative behaviours (biofilm formation, quorum-sensing, secretion systems, siderophores production and usage, and antibiotic degradation). We employed the KEGG Orthology (KO) database^37^ for functional annotations and used KofamScan^38^ to assign KO identifiers to protein sequences.

To create a list of “cooperative KO identifiers”, we first generated a list of cooperative keywords by referring to prior studies^36^. We then downloaded the complete KO identifier list ( https://www.genome.jp/ftp/db/kofam/ko_list.gz) and filtered it based on these keywords. This list was then manually curated to ensure specificity and relevance to prokaryotes, resulting in a list of 386 functionally annotated cooperative KOs (Table S4).

To compare our results with previous studies^19,20^, we also considered genes coding for extracellular proteins as putatively genes for cooperative traits. This is because extracellular proteins often act as public goods, whose benefits are shared with neighbouring cells^21,36^. We first downloaded a dataset of proteins whose subcellular localization (SCL) has been verified by laboratory experimentation^63^ (http://db.psort.org/static/downloads/Experimental-PSORTdb-v4.00.tsv). We then retrieved extracellular proteins from the dataset and employed KofamScan to assign significant KO identifiers to these proteins, resulting in a list of 193 “extracellular KO identifiers”. This list will be integrated into the previous list of functionally annotated cooperative KO identifiers, and will be used to mine genes for cooperative traits in further steps. The final list included 579 cooperative KO identifiers (extracellular factor, 193; biofilm, 84; quorum sensing, 23; secretion systems, 82; siderophores, 41; and antibiotic degradation, 156; Table S4).

After assigning KO identifiers to every gene in our dataset and identifying genes for cooperative traits, we calculated the proportion of genes for cooperative traits for each genome. In calculating the proportions, a potential issue arose when some genes lack confident functional annotations, which could influence the accuracy of the estimates. To address this, we directly used KOs to represent gene content, specifically considering only genes that could be matched with at least one significant KO identifier. As a result, the proportion of genes for cooperative traits in each genome was calculated as the ratio of the number of cooperative KOs to the total number of KOs. At this stage, 25,785 species had both niche breadth annotations and calculated proportions of genes for cooperation. Our primary analyses then focused on these species (Table S5).

### Phylogeny

The phylogenetic reference tree of bacterial species was downloaded from GTDB^32^ (release 207, https://data.gtdb.ecogenomic.org/releases/release207/207.0/bac120_r207.tree). To focus on the 25,785 species relevant to our study, we pruned this phylogenetic tree using the “*keep.tip*” function in the R package *ape* (version 5.7.1)^64^, resulting a final phylogenetic tree with 25,785 species.

### Ancestral state reconstruction for niche breadth and gene content

To infer causality and calculate gene gain/loss rates, we estimated ancestral niche breadth and gene content for each species’ representative genome. For niche breadth, extant species had three discrete states: generalist, intermediate, and specialist. We applied an empirical Bayesian method (the marginal posterior probability approximation (MPPA) method with the F81-like model) in PastML^65^ (version 1.9.15) to reconstruct ancestral states for niche breadths of each internal node in our species’ phylogeny (Figure E3). Of the 25,784 internal nodes, 9,235 were estimated as generalists, 3,994 as specialists, and 11,683 as intermediates. Ancestral states for 872 nodes remained undetermined due to uncertainty of the method (Figure E4).

For gene content, we used a similar approach to estimate the ancestral presence or absence of each ortholog group (OG) (n = 10,060) for every internal node (n = 25,784) in the phylogeny using species’ representative genomes (Figure E3). The extant species’ OG status was determined through KEGG Orthologs (KO). Consequently, the reconstructed “genome” for each internal node (ancestral species) consisted of all KOs marked as “present”. The proportion of genes for cooperative traits for each internal node was then calculated as the ratio of the number of cooperative KOs to the total number of KOs present in that node. These proportions ranged from approximately 0 to 0.07 (Figure E4b).

### Causality inference

To probe the causal relationship between cooperation and niche breadth evolution, we focused on the chronological order in which these traits evolved, inspired by a common causal inference framework, Granger causality^66,67^. Specifically, if events A and B are correlated and event A precedes event B, we would suggest that A “causes” B. For instance, if carrying higher proportion of genes for cooperative traits (Event A) is found to precede becoming a generalist (Event B), we could propose that a higher proportion of genes for cooperation may facilitate the transition to a broader niche breadth (Figure E3).

To determine the chronological order of these evolutionary events, we compared the states of ancestral nodes (parents) with their immediate descendants (children). For instance, to investigate whether possessing more genes for cooperation promotes the transition to being a generalist, we examined the proportion of genes for cooperation in specialist parents whose descendants broaden their niche versus those whose descendants remain specialists. If the former group displays a higher proportion of genes for cooperation, it would suggest that the increased presence of genes for cooperation likely preceded the transition to a more generalist habitat preference. In such a case, we could propose that having more genes for cooperation facilitates the evolution transition from specialists to generalists (Figure E3).

### Estimation of gene gain and loss rates

We analysed the rate of gene gains and losses over two different evolutionary scales: across-species (relative longer-term) and within-species (relative short-term). On the across-species (interspecific) scale, the rate of gains and losses for each KEGG Orthologs (KO) was calculated as the average rate of state transitions per phylogenetic branch — either presence (coded as 1) to absence (coded as 0) or vice versa^42^. Gene gains were indicated by 0-to-1 transitions, while gene losses were signified by 1-to-0 transitions (Figure E5). The state transition rates for 10,060 KOs were determined through ancestral state reconstruction using PastML^65^ (version 1.9.15). In calculating interspecific gene gain and loss rates, we used the representative genomes of each species to create a species—KO presence/absence table for reconstructing ancestral states. It‘s important to note that species may have different genomes, and using different genomes could lead to variations in the species—KO presence/absence tables^68^. However, a previous study has shown that if a KO was present in the representative genome of a species, it was likely to be present in nearly all other genomes of that species^42^. Therefore, using representative genomes to calculate these rates is considered reliable for our analyses.

To assess whether genes for cooperative traits experienced more frequent gains and losses compared to genes for private traits, we examined the difference in average rates of state transitions per branch between cooperative and non-cooperative KOs (Table S7). Additionally, the net gain or loss rates for each cooperative KO were estimated using the “*asr_mk_model*” function in the R package *castor* (version 1.7.10), employing an “*ARD (all rates different)*” rate model^69^ (Figure E5).

On the within-species (intraspecific) scale, direct estimation of gene gain or loss rates is unfeasible without strain-level phylogeny^70^. We thus used the extent to which a gene is categorized as accessory in a pangenome to serve as an indirect indicator of the intraspecific rate of gene gains or losses. This is because accessory genes are more prone to being gained or lost compared to core genes^47^. We reconstructed pangenomes for species that had at least 100 high-quality genomes in GTDB, resulted in a list comprising 171 species with 155,554 genomes (Figure E6, Table S9). The typical process of pangenome reconstruction can be broken down into three primary steps: 1) gene prediction and annotation, 2) homologous genes identification, 3) categorizing genes as core or accessory based on their presence or absence across different genomes^71,72^. In our study, we employed the protocol outlined in the KO database for gene prediction and ortholog identification^37^. KofamScan^38^ was used to assign KO identifiers to protein sequences, thereby grouping genes with identical KO identifiers into gene families. Consequently, the presence or absence profile of each gene family across the genomes of each species was represented by the presence or absence of each KO.

To assess whether genes for cooperative traits are more likely to be accessory genes in pangenomes, we introduced an index π to gauge the extent to which these genes are accessory. For each species, we randomly selected two strains to create a minimum pangenome based on these pairwise genomes. We then estimated Π, which represents the difference between the observed and expected proportions of accessory genes for cooperative traits. The Π is,

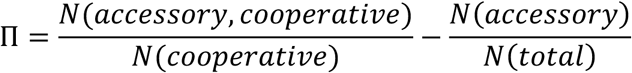

Here, *N*(*accessory*, *cooperative*) refers to the number of cooperative KOs that are accessory, *N*(*cooperative*) is the total number of cooperative KOs, *N*(*accessory*) is the number of accessory KOs, and *N*(*total*) is the total number of KOs in this minimum pangenome. This index allows us to quantify how much more (or less) likely a gene for cooperative traits is to be an accessory gene compared to what would be expected by chance in the minimum pangenome.

The index π is then denoted as the average Π calculated across all possible minimum pangenomes consisting of two genomes:

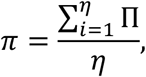

where 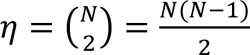, which is the total number of unique combinations of two genomes that can be chosen from *N* available strains for a species. The higher the value of π for genes for cooperative traits, the more they tend to be accessory genes within bacterial pangenomes, indicating a greater frequency of gains and losses.

For species with an extensive number of genomes, such as *Escherichia coli* which has 26,573 genomes in our dataset, the computation of Π for all selected genome pairs would be resource-intensive. To address this, we applied Central Limit Theorem (CLT) to estimate π for these species. This approach involved three steps: 1) randomly sample 20 genomes of a species for 10,000 times, 2) calculating the π for each sample, and 3) using the mean value of all the sample π as an estimation for π of the species. According to the CLT, the average of sample means will converge to the population mean when the sample size is sufficiently large.

### Diversification analysis

To explore the independent impact of cooperation or niche breadth on bacterial diversification, we utilized the binary-state speciation and extinction (BiSSE) model available in the R package *diversitree* (version 0.9.20) to fit our data^49^. The BiSSE model is used to assess the effect of binary traits (0 or 1) on speciation and extinction rates. We thus binarized two traits in our dataset: (i) the proportion of genes for cooperation, using the median proportion as a cut-off. Values below this cut-off were categorized as “lower” cooperation (0), and values above as “higher” cooperation (1; Figure E8a). (ii) For niche breadth, species found in less than 6 habitat clusters were classified as having “narrower” niche breadth (0), and those in 6 or more habitat clusters as having “broader” niche breadth (1), based on the distribution of niche breadths across all species (Figure E8d). The BiSSE model outputs six parameters for each input phylogenetic tree and trait data: two speciation rates (λ_0_ *and* λ_1_), two extinction rates (μ_0_ *and* μ_1_), and two state-transition rates (from state 1 to 0 as *q*_10_, and from state 0 to 1 as *q*_01_).

We identified two potential issues with fitting the BiSSE model and other character-dependent SSE models. First, using a phylogenetic tree with a large number of species can lead to implausible results, such as near-zero extinction rates (Figure E7). Second, fitting too many parameters can increase the risk of overfitting, complicating the interpretation of the results. To address these concerns, we implemented two strategies. First, instead of using a single phylogenetic tree for all species, we fitted the BiSSE models (and all other character-dependent SSE models) separately for each family containing at least 100 species (Table S10). This approach reduces the likelihood of errors associated with large tree sizes and allows for comparisons of diversification patterns across different taxa. Second, to simplify the model and further mitigate the risk of overfitting, we set the two state-transition rates to be equal (*q*_10_ = *q*_01_).

To explore the collective effect of cooperation and niche breadth on bacterial diversification, we integrated these two binary traits into a single trait with four states. We then applied the multi-state speciation and extinction (MuSSE) model, implemented in the *diversitree* package, to study the impact of this newly integrated trait on diversification. The four states are: 1 (narrower niche breadth and lower cooperation), 2 (narrower niche breadth and higher cooperation), 3 (broader niche breadth and lower cooperation), and 4 (broader niche breadth and higher cooperation). To simplify the MuSSE model, we set all state-transition rates equal to the rate between state 1 and 2 (*q*_12_). The model thus outputs 9 parameters: four speciation rates (λ_1_, λ_2_, λ_3_, λ_4_), four extinction rates (μ_1_, μ_2_, μ_3_, μ_4_), and one state-transition rate (*q*_12_).

The R package *diversitree* includes a built-in model specifically designed to investigate the impact of continuous traits on diversification, i.e., the quantitative-state speciation and extinction (QuaSSE) model^49^. While QuaSSE is theoretically more appropriate for examining the role of the proportion of genes for cooperation in shaping diversification, we found that implementing the QuaSSE model is impractical for complex phylogenetic trees with more than about 100 species. Consequently, we opted to use models for discrete traits, such as the BiSSE and MuSSE models, to examine the role of cooperation in diversification.

### Statistics

R package *MCMCglmm* (version 2.35) allows us to fit generalized linear mixed-effects models (GLMMs) to study the relationships between variables while controlling for phylogenetic relationships of species, using a Markov chain Monte Carlo approach with a Bayesian statistical framework^39^. To account for potential phylogenetic nonindependence among the species in our dataset, we incorporated the phylogeny as a random effect in the model. To ensure that the phylogenetic tree used in the MCMCglmm fitting had an ultrametric structure, we applied the “*force.ultrametric*” function in the R package “*phytools*” (with an option *“method = extend”*) to transfer our tree into an ultrametric form^73^. To assess the goodness of fit for each model, we reported the *pMCMC* value (referred to as “p-value”). When conducting causal inference, it’s necessary to control for phylogenetic relationships between ancestral species (nodes) rather than extant species (tips). This can be achieved by using the “*inverseA*” function in the R package “*MCMCglmm*” to invert phylogenetic covariance matrices that include nodes. For diversification analyses, the most recent common ancestor of all species within a family is identified as the family node. Thus, the phylogenetic relationships between families can be represented through the phylogenetic relationships of these family nodes. This approach allows the application of similar strategies used in causal inference to control for the relationships between families while performing diversification analyses.

## Supporting information

Supplementary materials (without tables)

## Acknowledgements

We thank Thomas Richards, Sira Sriswasdi, Anna Dewar, Zheren Zhang, Xiaotong Zhang and Jordi Bascompte for helpful discussions. We thank the National Science Foundation of China (31530007: C.H.), the Japan Society for the Promotion of Science (KAKENHI 22J20318: N.K.; KAKENHI 22H04925: W.I.), the Japan Science and Technology Agency (JPMJGX23B2, JPMJCR19S2: W.I.), European Research Council (834164: C.H., L.J.B. and S.A.W.) for funding.

## Contributions

Conceptualization: C.H., N.K. and S.A.W.; Data collection: C.H. and N.K.; Methodology: C.H., N.K., M.I., L.J.B., W.I. and S.A.W.; Investigation: C.H. and N.K.; Visualization: C.H. and N.K.; Funding acquisition: W.I. and S.A.W.; Supervision: W.I. and S.A.W.; Writing—original draft: C.H.; Writing—review and editing: C.H., N.K., M.I., W.I. and S.A.W.

## References

1. Lozupone, C. A. & Knight, R. Global patterns in bacterial diversity. Proceedings of the National Academy of Sciences 104, 11436–11440 (2007).

2. Thompson, L. R. et al. A communal catalogue reveals Earth’s multiscale microbial diversity. Nature 551, 457–463 (2017).

3. Delgado-Baquerizo, M. et al. A global atlas of the dominant bacteria found in soil. Science 359, 320–325 (2018).

4. Lima-Mendez, G. et al. Determinants of community structure in the global plankton interactome. Science 348, 1262073 (2015).

5. Wu, L. et al. Global diversity and biogeography of bacterial communities in wastewater treatment plants. Nat Microbiol 4, 1183–1195 (2019).

6. Sexton, J. P., McIntyre, P. J., Angert, A. L. & Rice, K. J. Evolution and Ecology of Species Range Limits. Annual Review of Ecology, Evolution, and Systematics 40, 415– 436 (2009).

7. Sexton, J. P., Montiel, J., Shay, J. E., Stephens, M. R. & Slatyer, R. A. Evolution of Ecological Niche Breadth. Annual Review of Ecology, Evolution, and Systematics 48, 183–206 (2017).

8. 8. Muller, E. E. L. Determining Microbial Niche Breadth in the Environment for Better Ecosystem Fate Predictions. mSystems 4, 10.1128/msystems.00080-19 (2019).

9. Sriswasdi, S., Yang, C. & Iwasaki, W. Generalist species drive microbial dispersion and evolution. Nat Commun 8, 1162 (2017).

10. von Meijenfeldt, F. A. B., Hogeweg, P. & Dutilh, B. E. A social niche breadth score reveals niche range strategies of generalists and specialists. Nat Ecol Evol 7, 768–781 (2023).

11. Hernandez, D. J., Kiesewetter, K. N., Almeida, B. K., Revillini, D. & Afkhami, M. E. Multidimensional specialization and generalization are pervasive in soil prokaryotes. Nat Ecol Evol 7, 1408–1418 (2023).

12. Carscadden, K. A. et al. Niche Breadth: Causes and Consequences for Ecology, Evolution, and Conservation. The Quarterly Review of Biology 95, 179–214 (2020).

13. Jaffe, A. L., Castelle, C. J. & Banfield, J. F. Habitat Transition in the Evolution of Bacteria and Archaea. Annual Review of Microbiology 77, null (2023).

14. Hall-Stoodley, L., Costerton, J. W. & Stoodley, P. Bacterial biofilms: from the Natural environment to infectious diseases. Nat Rev Microbiol 2, 95–108 (2004).

15. West, S. A., Griffin, A. S., Gardner, A. & Diggle, S. P. Social evolution theory for microorganisms. Nat Rev Microbiol 4, 597–607 (2006).

16. West, S. A. & Cooper, G. A. Division of labour in microorganisms: an evolutionary perspective. Nat Rev Microbiol 14, 716–723 (2016).

17. West, S. A., Cooper, G. A., Ghoul, M. B. & Griffin, A. S. Ten recent insights for our understanding of cooperation. Nat Ecol Evol 5, 419–430 (2021).

18. Griffin, A. S., West, S. A. & Buckling, A. Cooperation and competition in pathogenic bacteria. Nature 430, 1024–1027 (2004).

19. McNally, L., Viana, M. & Brown, S. P. Cooperative secretions facilitate host range expansion in bacteria. Nat Commun 5, 4594 (2014).

20. Garcia-Garcera, M. & Rocha, E. P. C. Community diversity and habitat structure shape the repertoire of extracellular proteins in bacteria. Nat Commun 11, 758 (2020).

21. Nogueira, T. et al. Horizontal Gene Transfer of the Secretome Drives the Evolution of Bacterial Cooperation and Virulence. Current Biology 19, 1683–1691 (2009).

22. Nogueira, T., Touchon, M. & Rocha, E. P. C. Rapid Evolution of the Sequences and Gene Repertoires of Secreted Proteins in Bacteria. PLOS ONE 7, e49403 (2012).

23. Dewar, A. E. et al. Plasmids do not consistently stabilize cooperation across bacteria but may promote broad pathogen host-range. Nat Ecol Evol 5, 1624–1636 (2021).

24. Hao, C., Dewar, A. E., West, S. A. & Ghoul, M. Gene transferability and sociality do not correlate with gene connectivity. Proceedings of the Royal Society B: Biological Sciences 289, 20221819 (2022).

25. Chen, Y.-J. et al. Metabolic flexibility allows bacterial habitat generalists to become dominant in a frequently disturbed ecosystem. ISME J 15, 2986–3004 (2021).

26. Di Rienzi, S. C. et al. The human gut and groundwater harbor non-photosynthetic bacteria belonging to a new candidate phylum sibling to Cyanobacteria. eLife 2, e01102 (2013).

27. Niehus, R., Mitri, S., Fletcher, A. G. & Foster, K. R. Migration and horizontal gene transfer divide microbial genomes into multiple niches. Nat Commun 6, 8924 (2015).

28. Dillard, J. R. & Westneat, D. F. Disentangling the Correlated Evolution of Monogamy and Cooperation. Trends in Ecology & Evolution 31, 503–513 (2016).

29. Cornwallis, C. K. et al. Cooperation facilitates the colonization of harsh environments. Nat Ecol Evol 1, 1–10 (2017).

30. Morlon, H. et al. Phylogenetic Insights into Diversification. (2024) doi:10.1146/annurev-ecolsys-102722-020508.

31. Xu, Q. et al. Microbial generalists and specialists differently contribute to the community diversity in farmland soils. Journal of Advanced Research 40, 17–27 (2022).

32. Parks, D. H. et al. GTDB: an ongoing census of bacterial and archaeal diversity through a phylogenetically consistent, rank normalized and complete genome-based taxonomy. Nucleic Acids Research 50, D785–D794 (2022).

33. Mise, K. & Iwasaki, W. Environmental Atlas of Prokaryotes Enables Powerful and Intuitive Habitat-Based Analysis of Community Structures. iScience 23, 101624 (2020).

34. Coelho, L. P. et al. Towards the biogeography of prokaryotic genes. Nature 601, 252–256 (2022).

35. Belcher, L. J. et al. SOCfinder: a genomic tool for identifying cooperative genes in bacteria. 2023.10.16.562460 Preprint at 10.1101/2023.10.16.562460 (2023).

36. Simonet, C. & McNally, L. Kin selection explains the evolution of cooperation in the gut microbiota. PNAS 118, (2021).

37. Kanehisa, M., Sato, Y., Kawashima, M., Furumichi, M. & Tanabe, M. KEGG as a reference resource for gene and protein annotation. Nucleic Acids Research 44, D457– D462 (2016).

38. Aramaki, T. et al. KofamKOALA: KEGG Ortholog assignment based on profile HMM and adaptive score threshold. Bioinformatics 36, 2251–2252 (2020).

39. Hadfield, J. D. MCMC Methods for Multi-Response Generalized Linear Mixed Models: The MCMCglmm R Package. Journal of Statistical Software 33, 1–22 (2010).

40. Domingo-Sananes, M. R. & McInerney, J. O. Mechanisms That Shape Microbial Pangenomes. Trends in Microbiology 29, 493–503 (2021).

41. Hall, R. J., et al. Gene-Gene Relationships in an Escherichia Coli Accessory Genome Are Linked to Function and Mobility. 2021.03.26.437181 https://www.biorxiv.org/content/10.1101/2021.03.26.437181v1 (2021) doi:10.1101/2021.03.26.437181.

42. Konno, N. & Iwasaki, W. Machine learning enables prediction of metabolic system evolution in bacteria. Science Advances 9, eadc9130 (2023).

43. Revell, L. J. & Harmon, L. J. Phylogenetic Comparative Methods in R. (Princeton University Press, 2022).

44. Kramer, J., Özkaya, Ö. & Kümmerli, R. Bacterial siderophores in community and host interactions. Nat Rev Microbiol 18, 152–163 (2020).

45. McCutcheon, J. P. & Moran, N. A. Extreme genome reduction in symbiotic bacteria. Nat Rev Microbiol 10, 13–26 (2012).

46. Rolland, J. et al. Conceptual and empirical bridges between micro- and macroevolution. Nat Ecol Evol 7, 1181–1193 (2023).

47. McInerney, J. O., McNally, A. & O’Connell, M. J. Why prokaryotes have pangenomes. Nat Microbiol 2, 1–5 (2017).

48. Maddison, W. P., Midford, P. E. & Otto, S. P. Estimating a Binary Character’s Effect on Speciation and Extinction. Systematic Biology 56, 701–710 (2007).

49. FitzJohn, R. G. Diversitree: comparative phylogenetic analyses of diversification in R. Methods in Ecology and Evolution 3, 1084–1092 (2012).

50. Maddison, W. P. & FitzJohn, R. G. The unsolved challenge to phylogenetic correlation tests for categorical characters. Syst Biol 64, 127–136 (2015).

51. Sun, S.-J. et al. Climate-mediated cooperation promotes niche expansion in burying beetles. eLife 3, e02440 (2014).

52. Duffy, J. E. & Macdonald, K. S. Kin structure, ecology and the evolution of social organization in shrimp: a comparative analysis. Proceedings of the Royal Society B: Biological Sciences 277, 575–584 (2009).

53. Rabosky, D. L. LIKELIHOOD METHODS FOR DETECTING TEMPORAL SHIFTS IN DIVERSIFICATION RATES. Evolution 60, 1152–1164 (2006).

54. Cornwallis, C. K. et al. Symbioses shape feeding niches and diversification across insects. Nat Ecol Evol 7, 1022–1044 (2023).

55. Bowers, R. M. et al. Minimum information about a single amplified genome (MISAG) and a metagenome-assembled genome (MIMAG) of bacteria and archaea. Nat Biotechnol 35, 725–731 (2017).

56. Benson, D. A. et al. GenBank. Nucleic Acids Research 41, D36–D42 (2013).

57. O’Leary, N. A. et al. Reference sequence (RefSeq) database at NCBI: current status, taxonomic expansion, and functional annotation. Nucleic Acids Research 44, D733–D745 (2016).

58. Quast, C. et al. The SILVA ribosomal RNA gene database project: improved data processing and web-based tools. Nucleic Acids Research 41, D590–D596 (2013).

59. Rognes, T., Flouri, T., Nichols, B., Quince, C. & Mahé, F. VSEARCH: a versatile open source tool for metagenomics. PeerJ 4, e2584 (2016).

60. Silby, M. W., Winstanley, C., Godfrey, S. A. C., Levy, S. B. & Jackson, R. W. Pseudomonas genomes: diverse and adaptable. FEMS Microbiology Reviews 35, 652–680 (2011).

61. Charrad, M., Ghazzali, N., Boiteau, V. & Niknafs, A. NbClust: An R Package for Determining the Relevant Number of Clusters in a Data Set. Journal of Statistical Software 61, 1–36 (2014).

62. A, K. Factoextra: extract and visualize the results of multivariate data analyses. R Package Version 1, (2016).

63. Lau, W. Y. V. et al. PSORTdb 4.0: expanded and redesigned bacterial and archaeal protein subcellular localization database incorporating new secondary localizations. Nucleic Acids Research 49, D803–D808 (2021).

64. Paradis, E. & Schliep, K. ape 5.0: an environment for modern phylogenetics and evolutionary analyses in R. Bioinformatics 35, 526–528 (2019).

65. Ishikawa, S. A., Zhukova, A., Iwasaki, W. & Gascuel, O. A Fast Likelihood Method to Reconstruct and Visualize Ancestral Scenarios. Molecular Biology and Evolution 36, 2069–2085 (2019).

66. Granger, C. W. J. Investigating Causal Relations by Econometric Models and Cross-spectral Methods. Econometrica 37, 424–438 (1969).

67. Detto, M. et al. Causality and Persistence in Ecological Systems: A Nonparametric Spectral Granger Causality Approach. The American Naturalist 179, 524–535 (2012).

68. Beavan, A. J. S., Domingo-Sananes, M. R. & McInerney, J. O. Contingency, repeatability, and predictability in the evolution of a prokaryotic pangenome. Proceedings of the National Academy of Sciences 121, e2304934120 (2024).

69. Louca, S. & Doebeli, M. Efficient comparative phylogenetics on large trees. Bioinformatics 34, 1053–1055 (2018).

70. Gamblin, J., Lambert, A. & Blanquart, F. Persistent, Private and Mobile genes: a model for gene dynamics in evolving pangenomes. 2024.07.15.603572 Preprint at 10.1101/2024.07.15.603572 (2024).

71. Page, A. J. et al. Roary: rapid large-scale prokaryote pan genome analysis. Bioinformatics 31, 3691–3693 (2015).

72. Tonkin-Hill, G. et al. Producing polished prokaryotic pangenomes with the Panaroo pipeline. Genome Biol 21, 180 (2020).

73. Revell, L. J. phytools: an R package for phylogenetic comparative biology (and other things). Methods in Ecology and Evolution 3, 217–223 (2012).

