## Supplementary materials (without tables) for "Cooperation shapes bacterial niche breadth evolution and patterns of diversification"

#### Section 1: Extended figures used to support findings in the main text.

**Figure E1.** The overall workflow of this study.

**Figure E2.** The distribution of niche breadths and proportions of genes for cooperative traits across species.

**Figure E3.** Inferring the causal relationship between cooperation and niche breadth evolution.

**Figure E4.** Reconstructed ancestral niche breadths and proportion of genes for cooperative traits.

**Figure E5.** Interspecific gene gain and loss rates of two cooperative KOs.

**Figure E6.** Phylogeny of species for pangenome reconstruction.

**Figure E7.** Challenges in fitting speciation and extinction (SSE) models using a large phylogenetic tree.

**Figure E8.** Diversification analyses using binary-state speciation and extinction (BiSSE) model.

**Figure E9.** Variation in speciation and extinction (SSE) model parameters across families.

**Figure E10.** Distribution of Silhouette index for clustering method "ward.D2".

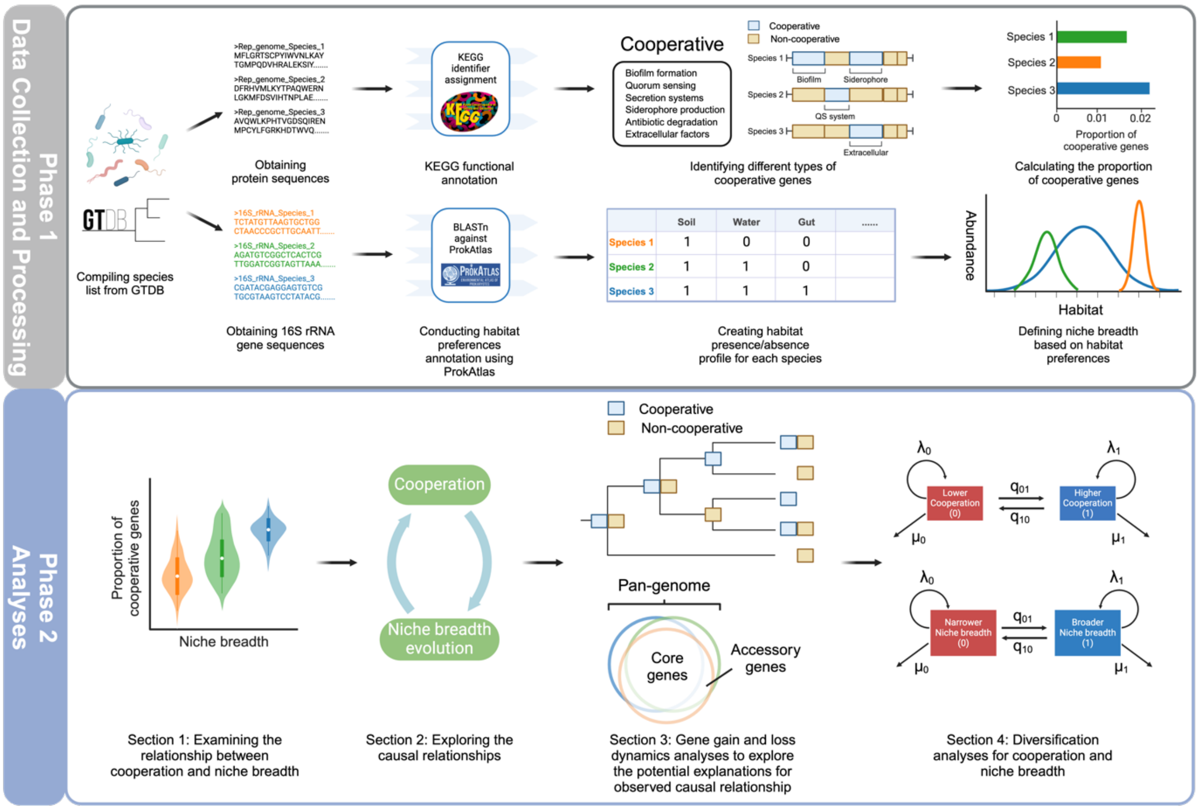


**Figure E1.** The overall workflow of this study. Phase 1: Data collection and processing. The bacterial species list was sourced from the Genome Taxonomy Database (GTDB). Protein sequences of representative genomes were downloaded from NCBI and analysed using the KEGG functional annotation tool to identify genes for cooperative traits. Genes were designated as "cooperative" if they were tagged with one or more of the following behaviours: biofilm formation, quorum sensing, secretion systems, siderophore production and usage, antibiotic degradation, and extracellular factor production. The proportion of genes for cooperation in each species' representative genome indicated their level of cooperative gene carriage. Subsequently, the 16S rRNA gene sequences from representative genomes were used to infer habitat preferences of each species using ProkAtlas, which includes metagenome-derived 16S rRNA gene sequences from 114 distinct habitats. These habitats were then grouped into 26 habitat clusters. Niche breadth for each species was quantified by the number of habitat clusters they occupied. Phase 2: Main analyses. The analyses consisted of four major sections. In section 1, we examined the relationship between cooperative gene carriage and niche breadth evolution. In section 2, we employed phylogenetic causal inference techniques to explore the causal relationship between cooperative gene carriage and niche breadth evolution. Section 3 involved analysing gene gain and loss dynamics to elucidate potential explanations for the observed causal patterns. At the interspecific level, gene gain and loss rates were measured by the rate of state transitions for each orthologous group (upper panel); at the intraspecific level, the rates of cooperative gene gain and loss are indicated by the propensity of a given cooperative gene to be an accessory gene in a pangenome, meaning these genes are present only in a subset of strains of a species (lower panel). Finally, in section 4, we analysed the role of cooperation and niche breadth in shaping bacterial diversification by fitting trait-dependent speciation ($\lambda$) and extinction ($\mu$) models, with different states of traits represented by different boxes.

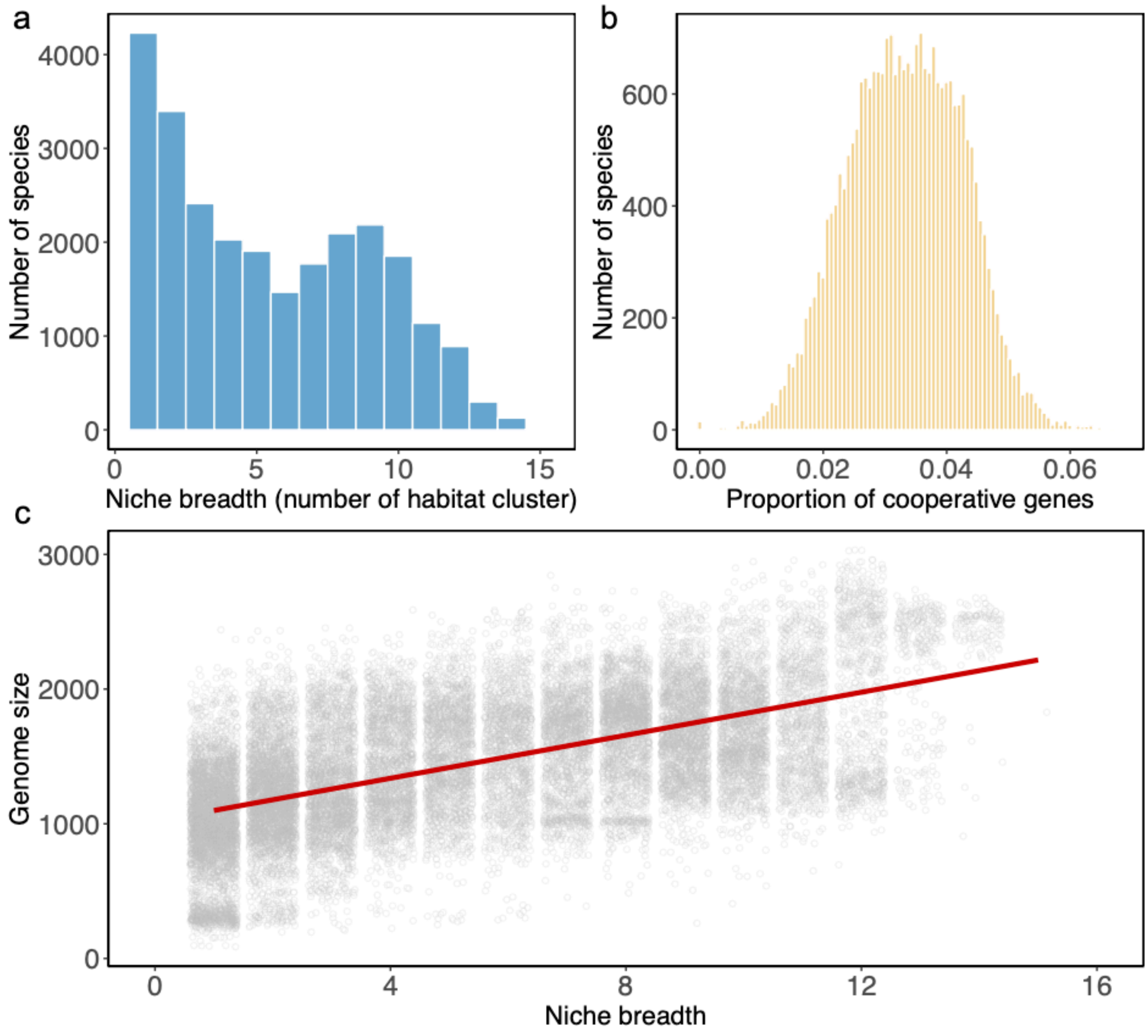


**Figure E2.** The distribution of niche breadths and proportions of genes for cooperative traits across species. (a) The distribution of niche breadths across species. The niche breadth of each species was quantified by the number of habitat clusters in which it was found. (b) The distribution of proportions of genes for cooperative traits across species. The proportion of genes for cooperative traits ranged from around 0 to 0.07. (c) The relationship between genome size and niche breadth. Species with broader niche breadth tend to have larger genome sizes.

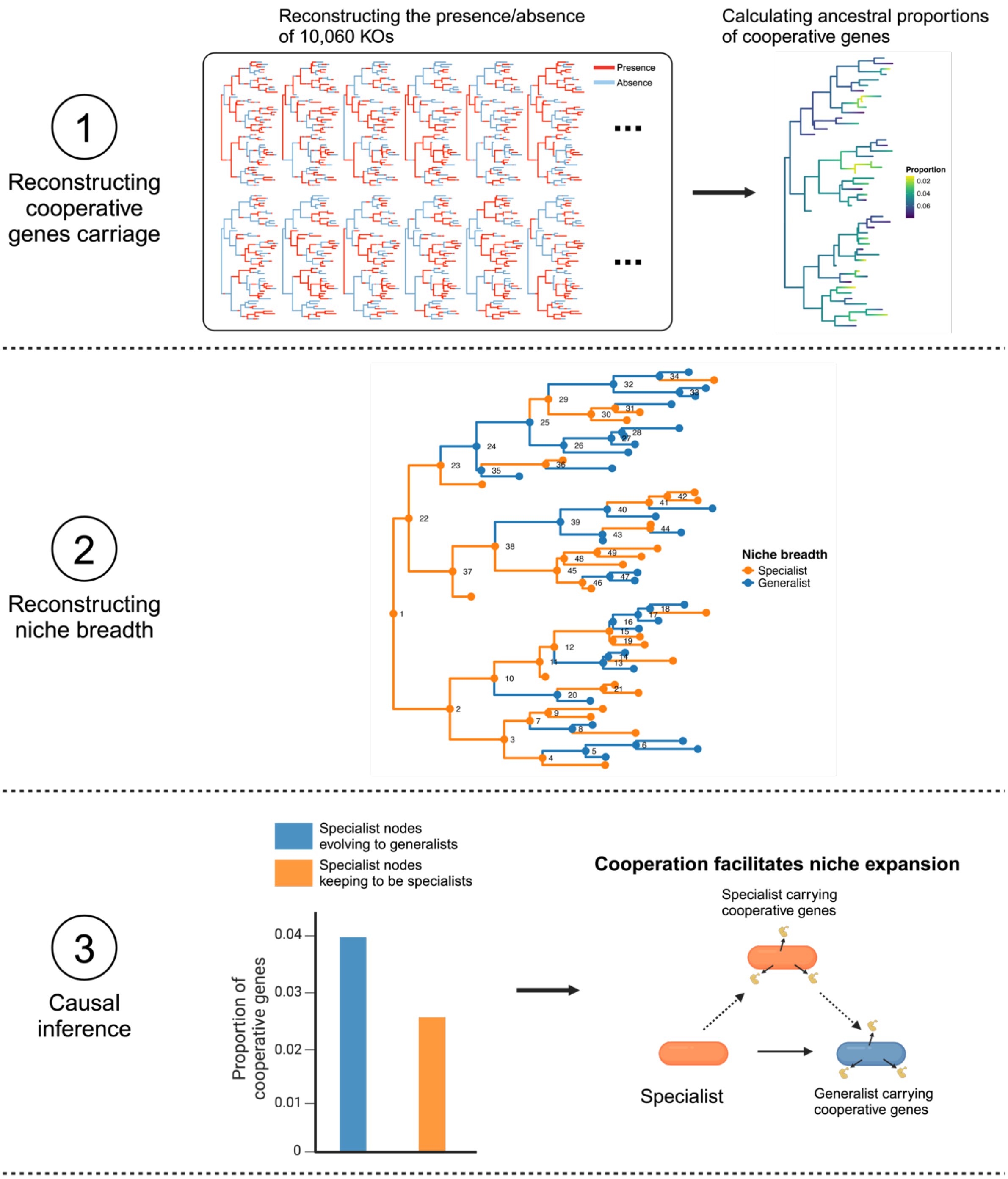


**Figure E3.** Inferring the causal relationship between cooperation and niche breadth evolution. To infer the causal relationship between cooperation and niche breadth evolution, we reconstructed the gene content and niche breadth of species in our dataset. For gene content, we reconstructed the ancestral presence or absence of each KEGG-defined orthologous group (KO; n = 10,060) for every internal node in the phylogeny. We then calculated the ancestral proportions of cooperative genes for each node by determining the ratio of the number of cooperative KOs to the total number of KOs present in that node. Next, we reconstructed the ancestral states for niche breadths of each internal node in the species' phylogeny. According to the hypothesis that cooperation facilitates niche expansion, we would expect that an increase in cooperative gene carriage precedes transitions from specialist to generalist states. To test this hypothesis, we compared the proportion of cooperative genes in specialist parent nodes whose descendants expanded their niche (e.g., nodes 4, 7, and 10) versus those whose descendants remained specialists (e.g., nodes 1, 2, and 3). If the former group displays a higher proportion of genes for cooperation, it will support the hypothesis that cooperation facilitates niche expansion. This technique can be applied to examine various evolutionary scenarios as outlined in Figure 4 (a-d).

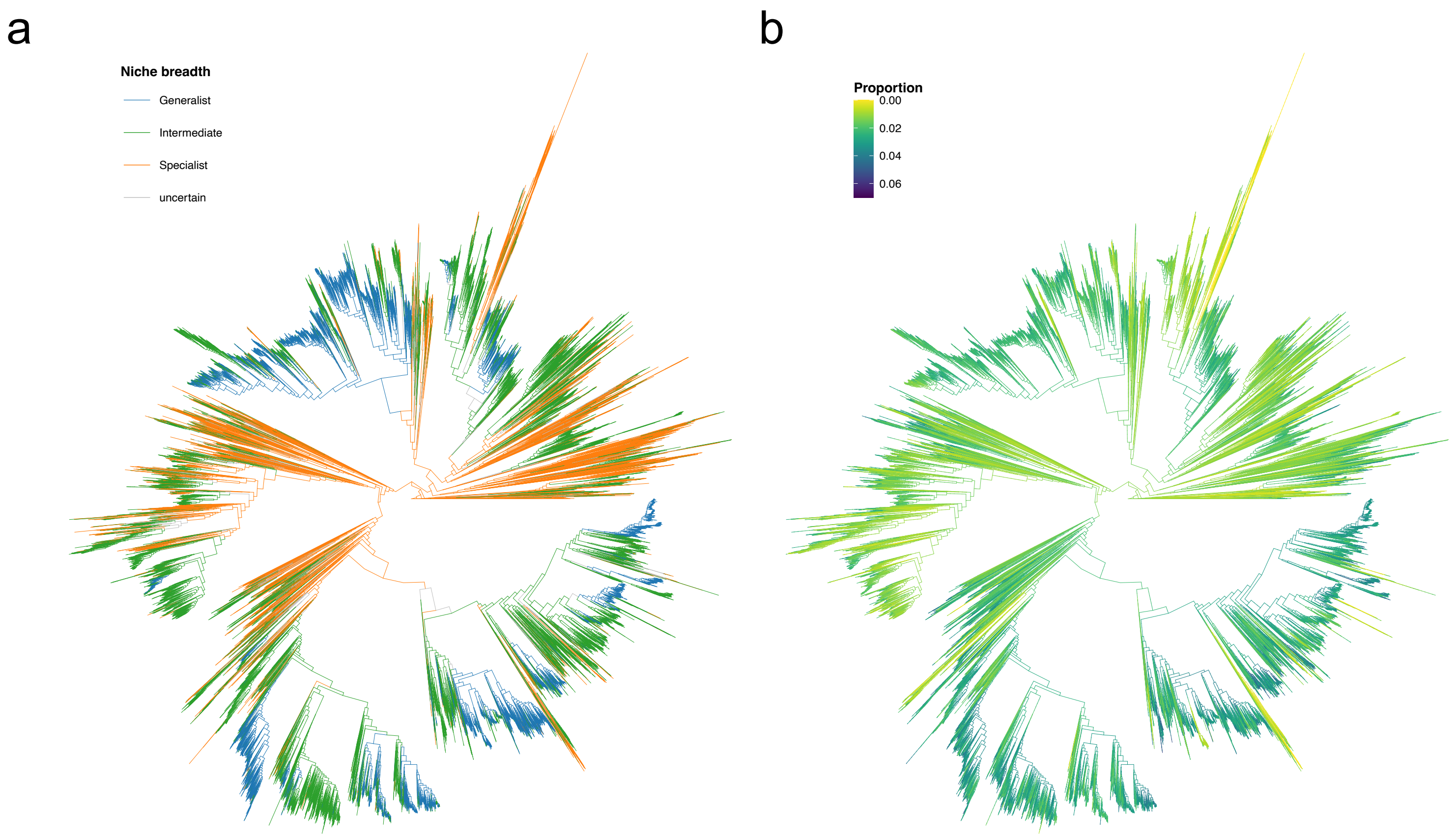


**Figure E4.** Reconstructed ancestral niche breadths and proportion of genes for cooperative traits. (a) We reconstructed niche breadths for 24,912 ancestral nodes, classifying them into 3,994 specialists (orange), 11,683 species with intermediate niche breadth (green), and 9,235 generalists (blue). The niche breadths for 872 nodes remained undetermined (grey). (b) Gene content profiles were reconstructed for 25,784 ancestral nodes. From these profiles, we calculated the ancestral proportion of genes for cooperation, which ranged from approximately 0 to 0.07. Darker branches have higher proportions of genes for cooperation, while lighter branches have lower proportions.

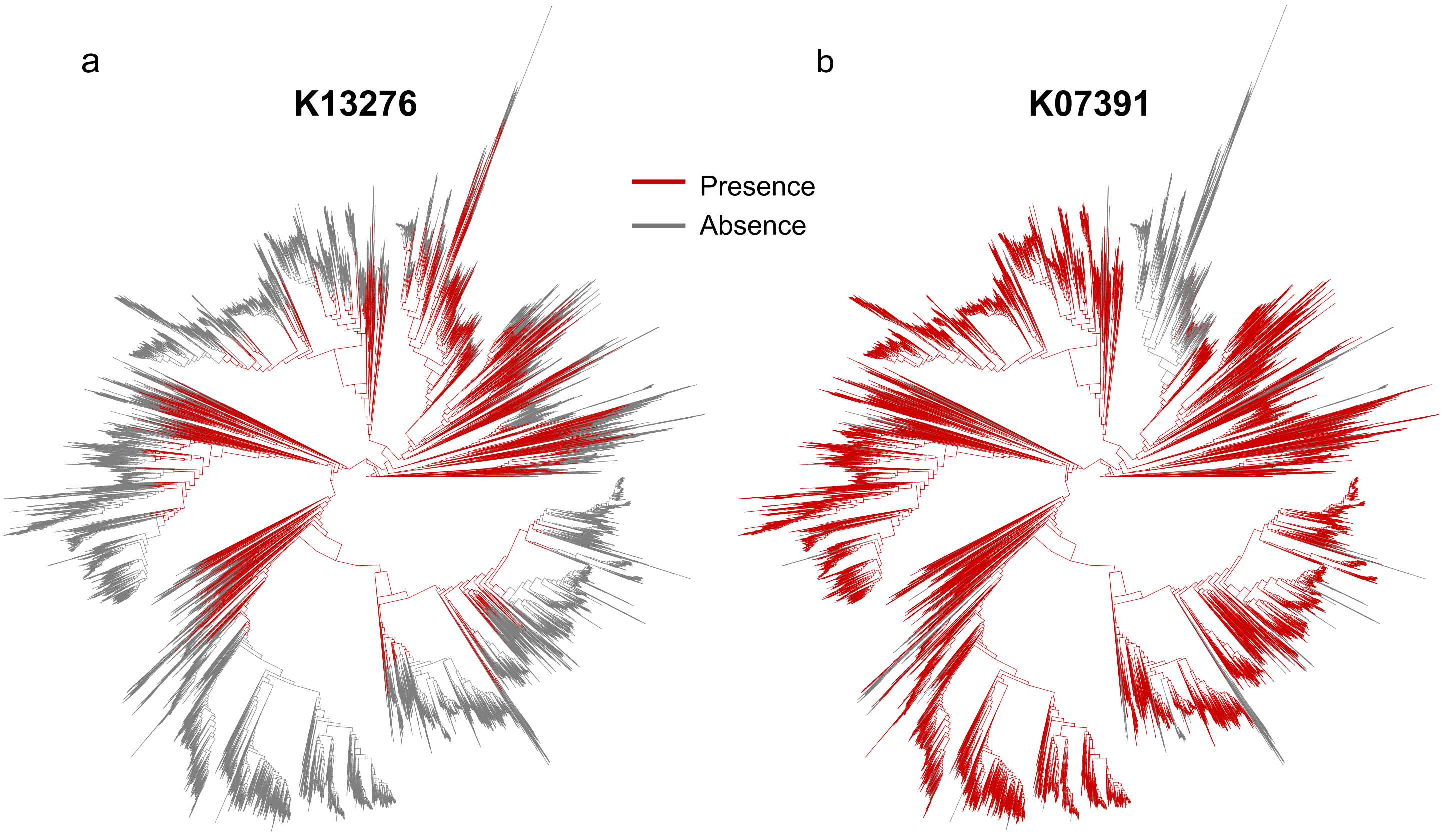


**Figure E5.** Interspecific gene gain and loss rates of two cooperative KOs (K13276 and K07391). The interspecific gain and loss rates of each KO were estimated by calculating the rate of state transitions using ancestral state reconstruction. Gene gains were indicated by transitions from absence to presence, while gene losses were denoted by transitions from presence to absence. Branches that possess a particular KO were coloured red, while branches without the KO were coloured grey. (a) The presence/absence profile of K13276 across the phylogenetic tree showed extensive, repeated gene loss, indicating that this KO underwent significant gene loss events. (b) The presence/absence profile of K07391 indicated a different pattern. K07391 did not experience significant gene loss; instead, its gain rate was higher than its loss rate.

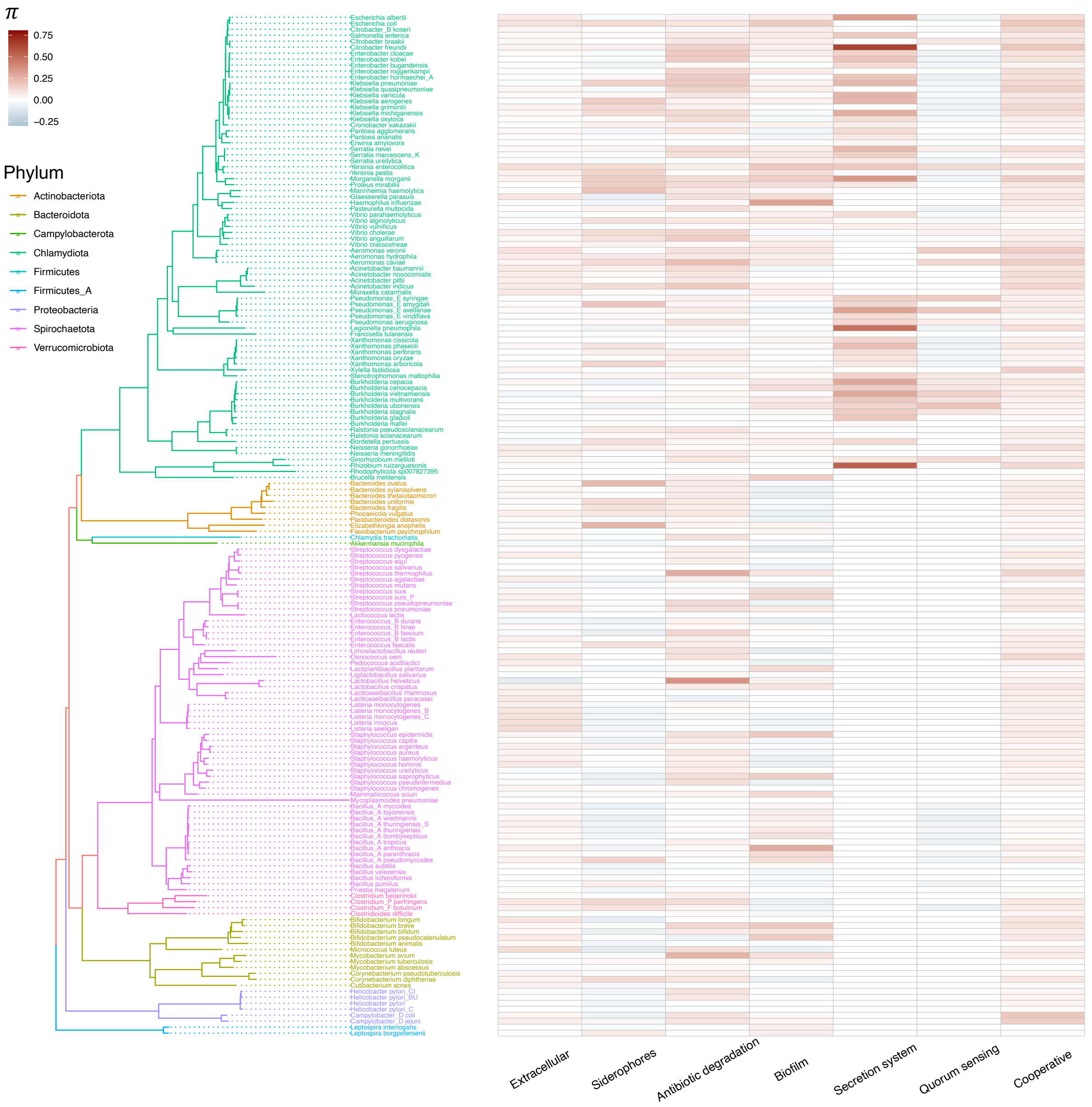


**Figure E6.** Phylogeny of species for pangenome reconstruction. 171 species were selected from the Genome Taxonomy Database (GTDB) for pangenome reconstruction. These species, each having at least 100 high-quality genomes in GTDB, belong to 9 distinct phyla. The branches of the phylogenetic tree are coloured to represent the different phyla to which these species belong. The heatmaps adjacent to the phylogenetic tree illustrate the $\pi$ values of genes coding various cooperative functions for every species, which indicates the propensity of genes for cooperation to be accessory genes within the pangenomes. A higher $\pi$ value suggests that genes for cooperation are more likely to be accessory, meaning they are present in only a subset of strains within a species. Blank grids in the heatmaps signify that a species does not possess a particular type of gene for cooperative traits.

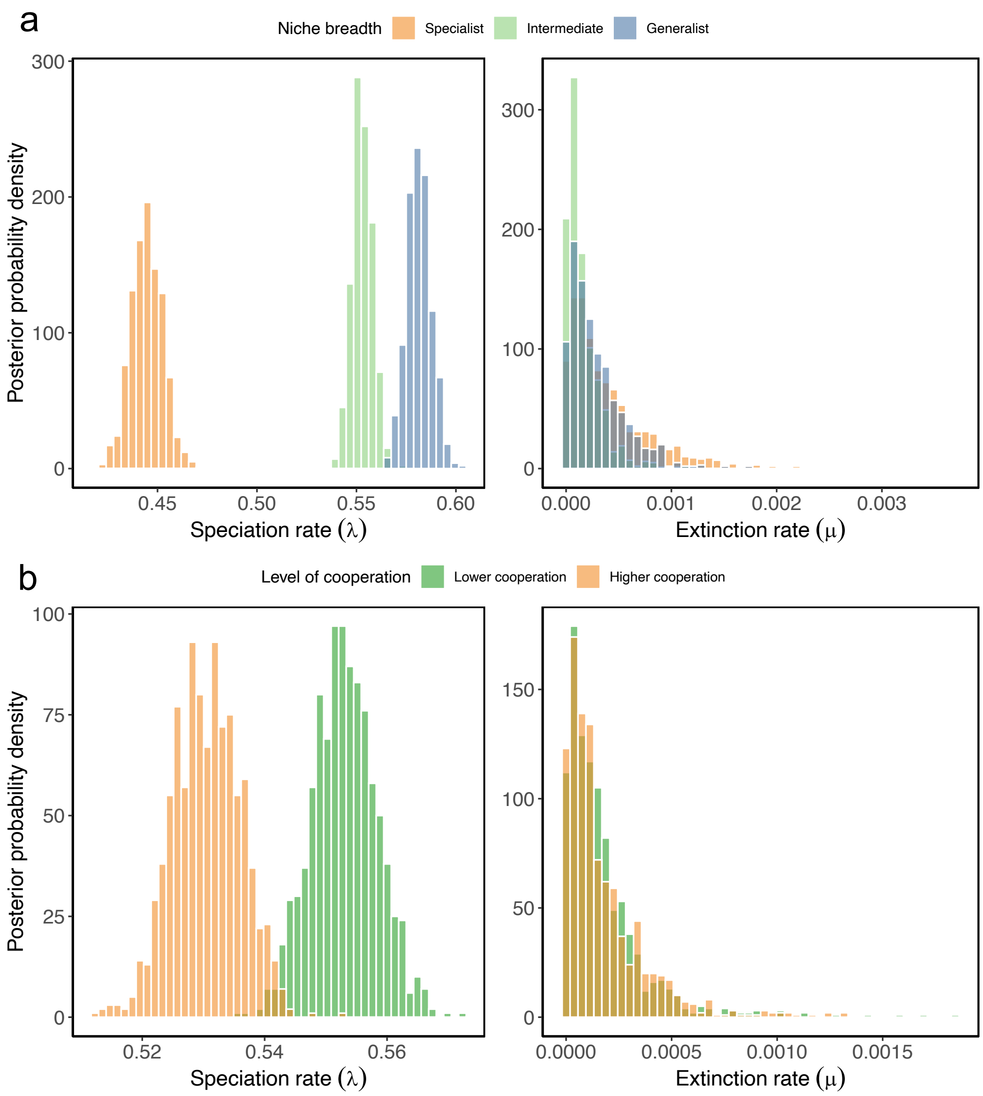


**Figure E7.** Challenges in fitting speciation and extinction (SSE) models using a large phylogenetic tree. We fitted two illustrative models to investigate the role of niche breadth and cooperation in bacterial diversification using a comprehensive phylogenetic tree comprising all species in our dataset (n = 25,785). The MuSSE model was used to examine niche breadth (with 3 states), and the BiSSE model was used to examine cooperation's impact on diversification. Bayesian analyses using Markov chain Monte Carlo (MCMC) methods were conducted to generate the posterior distributions of the parameters in these models, with 5,000 iterations. Panel (a) displays the posterior probability distributions for the parameters of MuSSE model related to niche breadth, while panel (b) displays the distributions for the parameters of BiSSE model related to cooperation. Both models predicted extremely low extinction rates. Consequently, to enhance the accuracy of model fittings and facilitate the comparison of diversification patterns across families, we fitted the SSE models for each family containing at least 100 species.

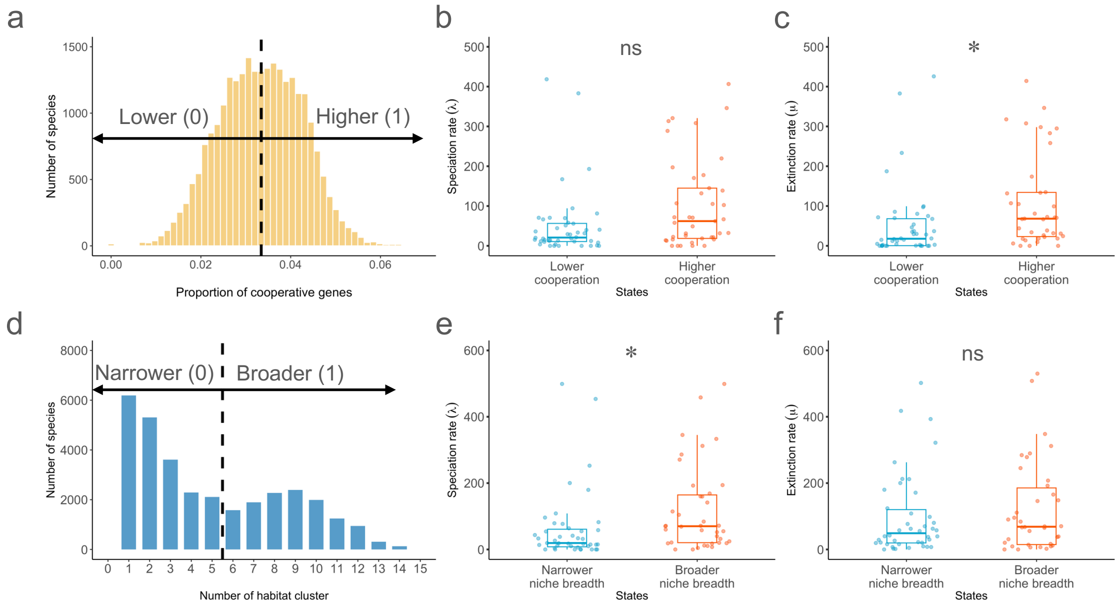


**Figure E8.** Diversification analyses using binary-state speciation and extinction (BiSSE) model. (a) The proportion of genes for cooperation was binarized using the median proportion as a cut-off. Values below this cut-off were categorized as "lower" cooperation (0), and values above as "higher" cooperation (1). (b) Levels of cooperation, whether lower or higher, do not influence speciation rates across families. Each dot represents a rate parameter for a given family. (c) Species with lower cooperation levels exhibited lower extinction rates compared to those with higher cooperation levels. (d) For niche breadth, species found in fewer than 6 habitat clusters were categorized as having "narrower" niche breadth (0), and those in 6 or more habitat clusters as having "broader" niche breadth (1). (e) Species with narrower niche breadth showed lower speciation rates than those with broader niche breadth. (f) Niche breadth, whether narrower or broader, does not impact extinction rates.

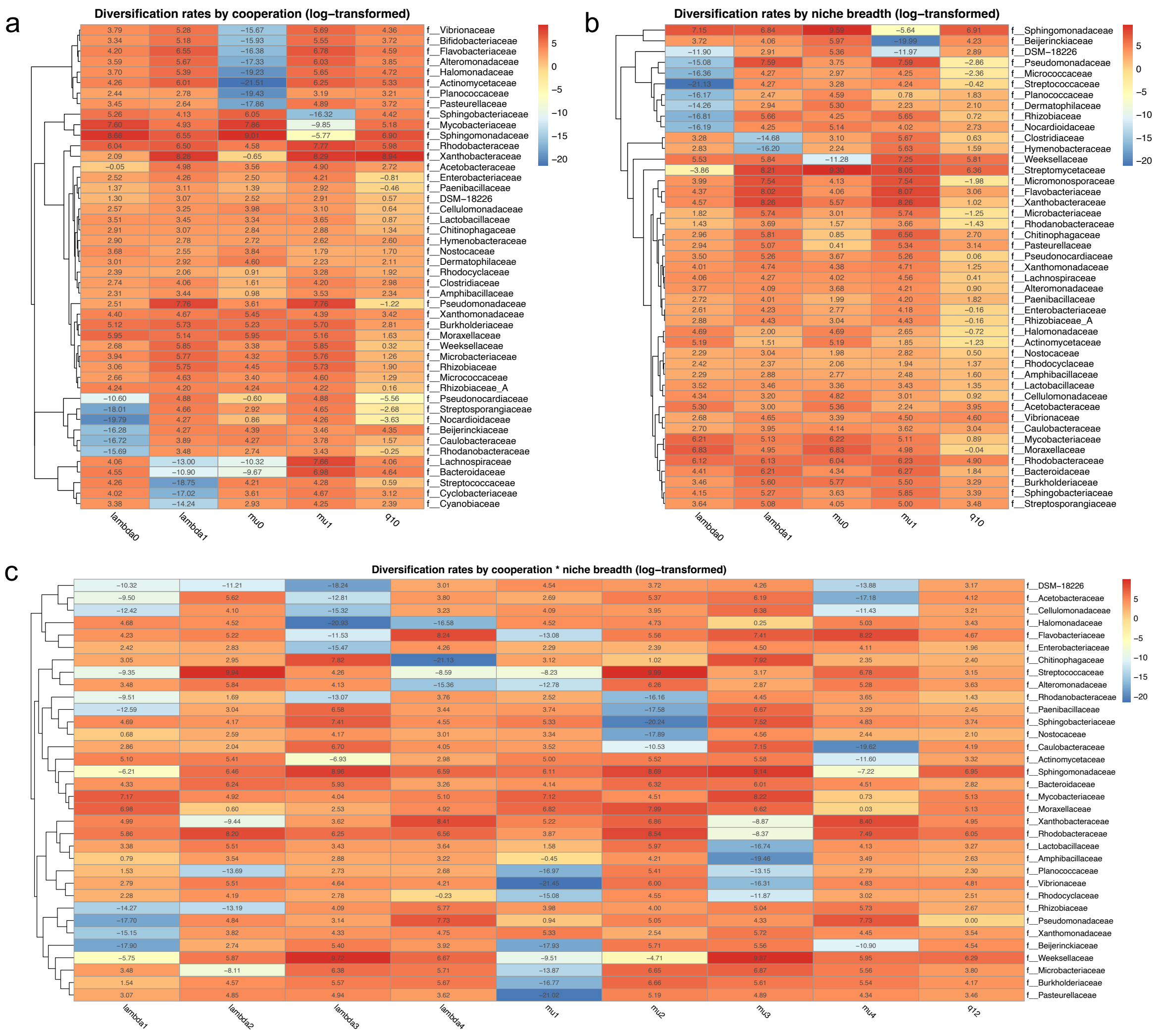


**Figure E9:** Variation in speciation and extinction (SSE) model parameters across families. All parameters were log-transformed to enhance visualization. (a) The heatmap illustrates the parameters of the BiSSE model used to study the impact of cooperation on bacterial diversification. In this context, the “0” state represents a lower level of cooperation, while the “1” state indicates a higher level of cooperation. For example, *lambda0* represents the speciation rates of species with lower cooperation, *mu1* represents the extinction rates of species with higher cooperation, and *q10* represents the transition rates between lower and higher cooperation states. (b) This heatmap illustrates the parameters of the BiSSE model focusing on niche breadth in bacterial diversification. The “0” state corresponds to a narrower niche breadth, whereas the “1” state represents a broader niche breadth. (c) The heatmap in this panel displays the parameters of the MuSSE model, which examines the collective effect of cooperation and niche breadth on bacterial diversification. The definitions of the states in this analysis are consistent with those used in Figure 6. The variation in these parameters highlights the diverse evolutionary strategies that may be employed by different bacterial families in response to cooperation and niche breadth evolution, influencing their speciation and extinction rates.

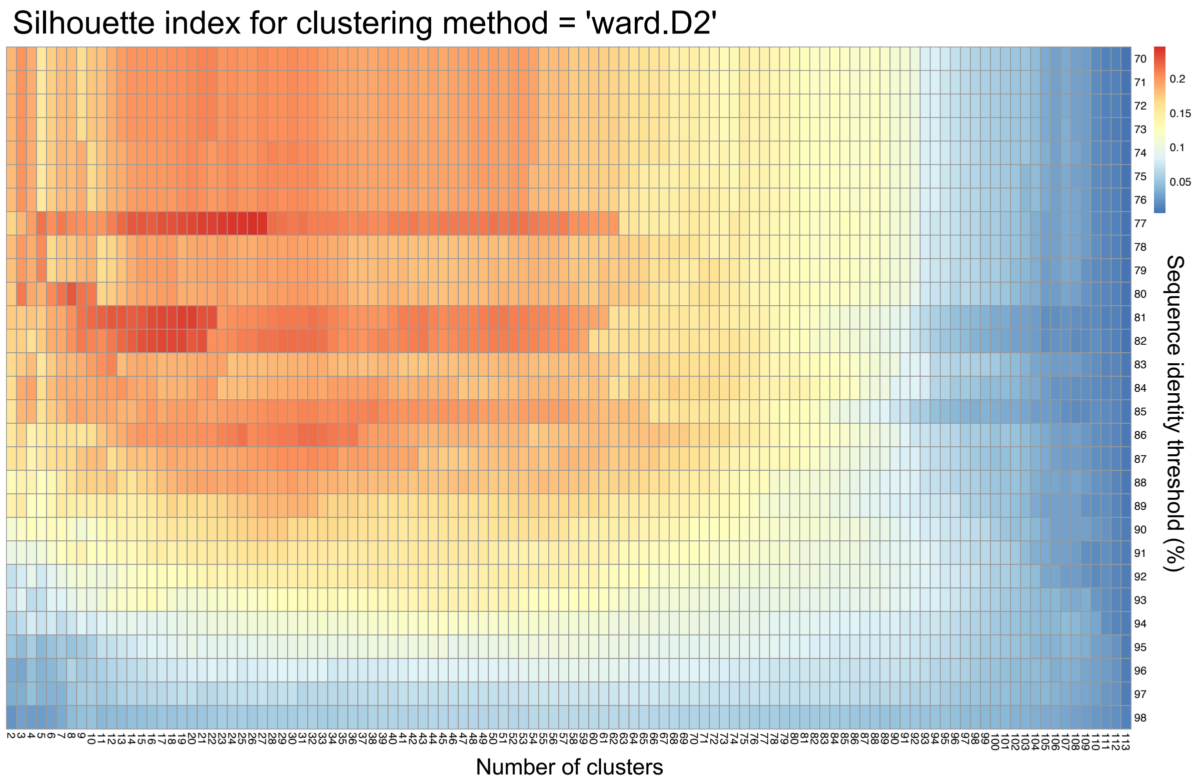


**Figure E10:** Distribution of Silhouette index for clustering method "ward.D2". To objectively determine the optimal clustering method, the number of clusters, and the sequence identity threshold $T$ for grouping ProkAtlas habitats into habitat clusters, we calculated the Silhouette index for each scenario. The Silhouette index, which ranges from −1 to +1, measures clustering performance, with higher values indicating better clustering. The heatmap displays the distribution of the Silhouette index across various sequence identity thresholds ($T$, y-axis) and numbers of clusters (x-axis) using the "ward.D2" clustering method. The optimal Silhouette index was achieved with $T$ = 77% and numbers of clusters = 26, which were then used for grouping ProkAtlas habitats into habitats clusters and defining species' niche breadths for subsequent analyses. The second optimal Silhouette index occurred at $T$ = 81% and numbers of clusters = 20. This alternative configuration was employed to test the robustness of the findings based on the optimal Silhouette index configuration (Supplementary section 2, part 1).

### Section 2: Robustness tests on the findings in the main text.

#### Part 1: Robustness tests on the relationship between cooperation and niche breadth evolution.

**Test 1:** *Species with broader niche breadths carried more genes for cooperative traits.*

**Test 2:** *The positive correlation between the proportion of genes for cooperation and niche breadth remained robust when using an alternative definition of niche breadth.*

**Test 3:** *The positive correlation between the proportion of genes for cooperation and niche breadth remained robust when using the second optimal threshold for clustering habitats.*

**Test 4:** *Genes for cooperation played a major role in niche breadth evolution among co-evolved genes.*

#### Part 2: Robustness tests on the patterns of causal inference.

**Test 1:** *Transition rates analysis confirmed the revealed direction of causality.*

**Test 2:** *Causal inference using the second optimal threshold for habitat clustering and niche breadth definition.*

#### Part 3: Robustness tests on the patterns of bacterial diversification.

**Test 1:** *The net diversification rates at the family level did not vary significantly across different states.*

#### Part 1: Robustness tests on the relationship between cooperation and niche breadth evolution

***Test 1:*** *Species with broader niche breadths carried more genes for cooperative traits.*

In the main text, we identified a positive correlation between the proportion of genes for cooperative traits and niche breadth across species. To verify the robustness of this finding, we further examined the relationship between the number of genes for cooperative traits and niche breadth across species. Our analysis confirmed that the number of such genes is positively correlated with niche breadth (MCMCglmm; n = 25,785 species; pMCMC < 0.001; Figure S1a, b; Table S1). This pattern persisted across different types of genes for cooperative traits, including those involved in antibiotic degradation, biofilm formation, quorum sensing, secretion systems, siderophore production, and extracellular protein production (MCMCglmm; n = 25,785 species; pMCMC < 0.001 for all categories; Figure S1b, c; Table S1).

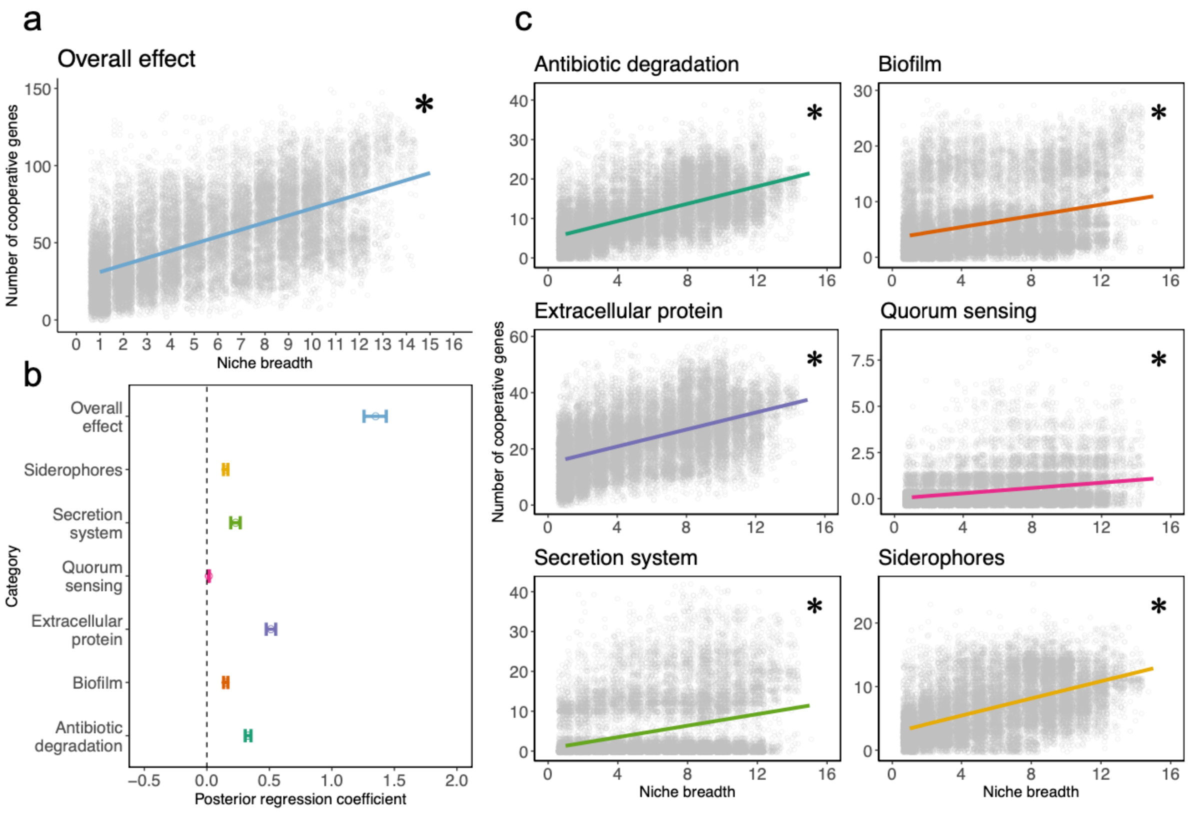

**Figure S1.** Correlation between number of genes for cooperation and niche breadth. (a) We found a positive correlation between the number of genes for cooperation and niche breadth. The asterisk denotes significant correlation. (b) The mean values and credible intervals of posterior regression coefficients for this correlation were derived from MCMCglmm model results. The dot represents the mean, and the horizontal bar represents the 95% credible interval of the posterior distribution of coefficients for each MCMCglmm model. (c) This positive correlation between the number of genes for cooperation and bacterial niche breadth was consistently observed across various functional categories of genes for cooperation. Asterisks denote significant correlations.

***Test 2:*** *The positive correlation between the proportion of genes for cooperation and niche breadth remained robust when using an alternative definition of niche breadth.*

The definition of niche breadth used in the main analysis was based on the number of habitat clusters occupied by species. In this test aiming to verify the findings from the main analysis, we redefined niche breadth by categorizing species into specialists, species with intermediate niche breadth, and generalists, similar to what we did when conducting causal inference in the main text (Table S5). We observed that habitat generalists carried a significantly higher proportion of genes for cooperation compared to both specialists and species with intermediate niche breadth (MCMCglmm; $n = 25{,}785$ species; generalist vs. intermediate species: pMCMC $< 0.001$; generalist vs. specialist: pMCMC $< 0.001$; Figure S2a; Table S1). Moreover, this pattern was consistent across different types of genes for cooperation (MCMCglmm; $n = 25{,}785$ species; pMCMC $< 0.001$ for all categories; Figure S2b; Table S1).

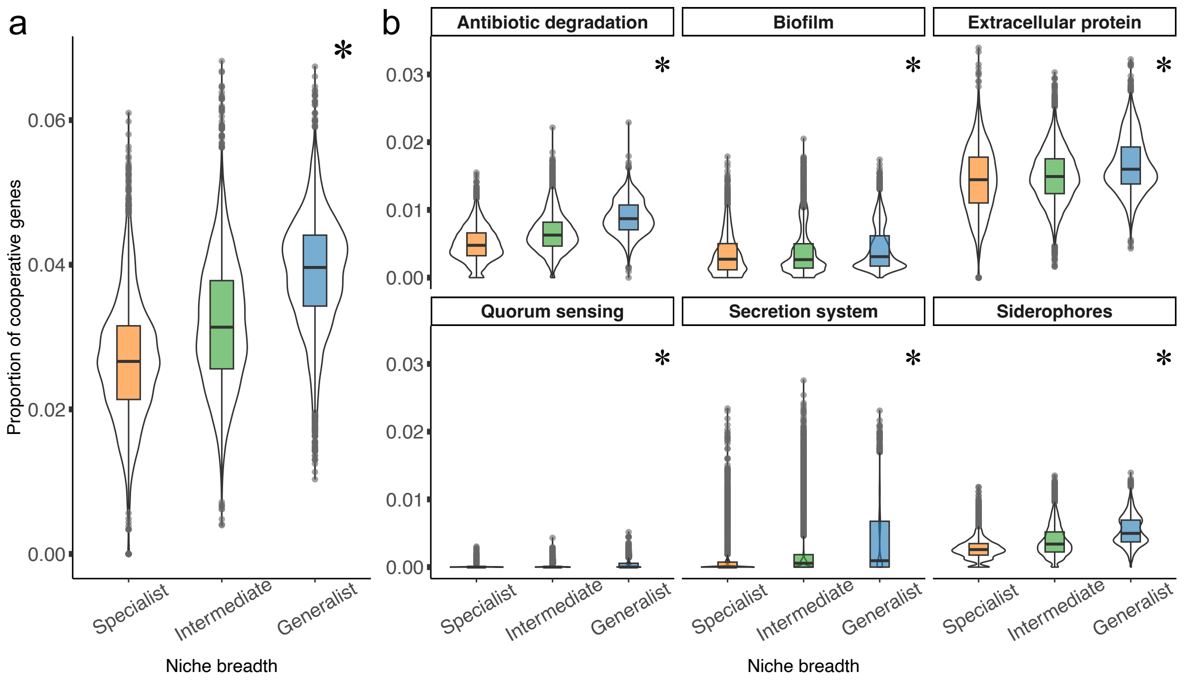


**Figure S2.** Correlation between cooperative gene proportion and bacterial niche breadth. (a) Habitat generalists consistently exhibited a higher proportion of cooperative genes compared to both specialists and species classified as intermediate. (b) Across varying functional categories of genes for cooperation, a consistent positive correlation between the proportion of these genes and bacterial niche breadth was observed. Asterisks denote significant correlations.

***Test 3:*** *The positive correlation between the proportion of genes for cooperation and niche breadth remained robust when using the second optimal threshold for clustering habitats.*

In the main analyses, we grouped ProkAtlas habitats using the "ward.D2" method with a sequence identity threshold $T$ = 77% and the number of clusters = 26. To verify the consistency of our findings under different clustering thresholds, we performed robustness tests using the second optimal threshold ($T$ = 81%, the number of clusters = 20) for habitat clustering (Figure S3a, Figure E10; Table S15). Niche breadths were then redefined based on the number of newly defined habitat clusters occupied by each species (Figure S3b; Table S15).

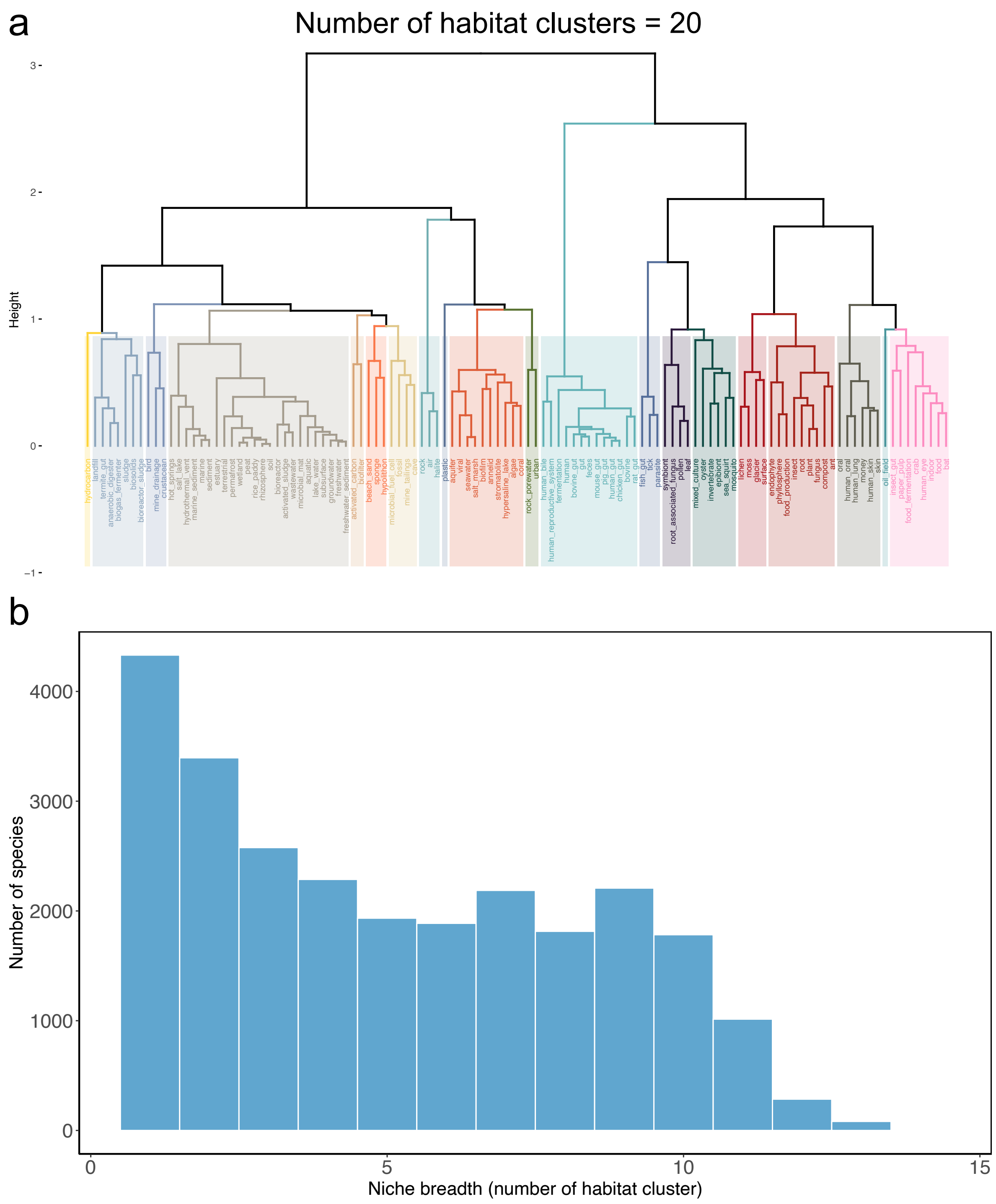


**Figure S3.** Assessment of bacterial niche breadths using the second optimal threshold for clustering habitats. The second optimal Silhouette index was identified at $T$ = 81% and numbers of clusters = 20. (a) 114 ProkAtlas habitats were grouped into 20 "habitat clusters" based on similarities in species composition. Species' habitat preference annotations were then updated accordingly. (b) The distribution of newly defined niche breadths across species. Niche breadth for each species was quantified by the number of habitat clusters in which they were present. Species found in more habitat clusters were considered to have broader niche breadths than those found in fewer clusters.

Based on the newly defined niche breadth, we re-examined the relationship between the proportion of genes for cooperative traits and niche breadth across species. The analysis confirmed that the positive correlation between the proportion of genes for cooperative traits and niche breadth was robust even with the updated definition of niche breadth (MCMCglmm; n = 25,785 species; pMCMC < 0.001; Figure S4a, b; Table S1). Furthermore, this consistent positive correlation was observed across various categories of genes for cooperation (MCMCglmm; n = 25,785 species; pMCMC < 0.001 for all categories; Figure S4b, c; Table S1).

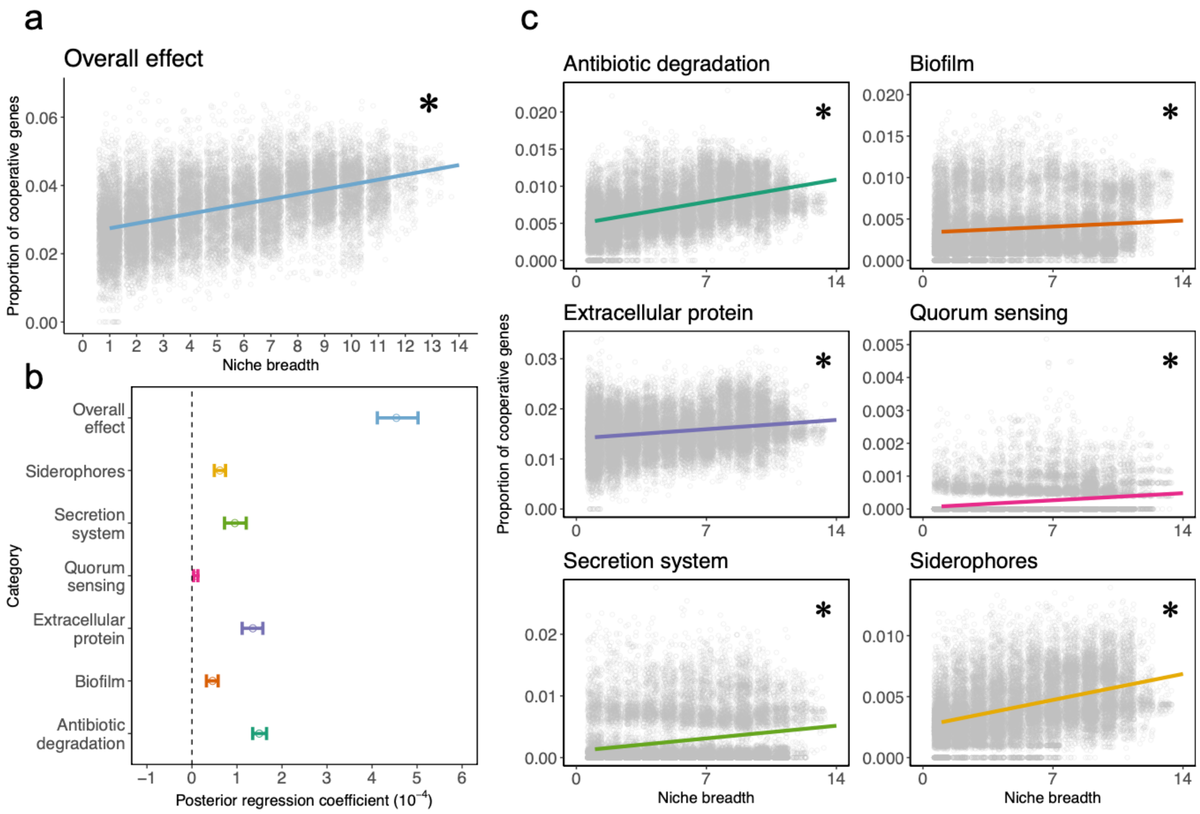

**Figure S4.** Correlation between proportion of genes for cooperation and newly defined niche breadth. (a) We observed a positive correlation between the proportion of cooperative genes and the newly defined niche breadth across species. (b) The mean values and credible intervals of the posterior regression coefficients for this correlation were derived from MCMCglmm model results. In the figure, the dot represents the mean value, while the horizontal bar indicates the 95% credible interval of the posterior distribution of the coefficients for each MCMCglmm model. (c) This positive correlation was consistently observed across various functional categories of cooperative genes.

***Test 4:*** *Genes for cooperation played a major role in niche breadth evolution among co-evolved genes.*

Notably, bacterial genes often experience simultaneous gains or losses, which is typically driven by coevolution among genes and associated with adaptation to diverse environments[1–3]. Specifically, genes within the same metabolic pathway are prone to coevolution due to their functional interdependencies[4–6]. In cases where one gene is lost or gained, it can render the entire pathway ineffective, potentially impacting another gene's function. This is particularly pertinent for genes for cooperative traits: if a gene for private traits that is functionally associated with a gene for cooperative traits also undergoes simultaneous gain or loss, gain or loss of this gene could confound the role of the gene for cooperation in driving niche breadth evolution. We thus examined the effect of these co-evolved genes.

<u>a) Identification of genes that co-evolved with genes for cooperation</u>

To pinpoint genes that may have co-evolved with genes for cooperation, we used the PhyloCorrelate tool to identify KEGG-defined orthologous groups (KOs) that share similar phylogenetic distribution patterns with cooperative KOs throughout the tree of life[7]. For each cooperative KO, we calculated the runs-adjusted Jaccard coefficient (rJC) between its presence/absence pattern (phylogenetic profile) and those of all other KOs, to quantify the similarities of all pairs of patterns across the phylogenetic tree. The 'runs-adjustment' method served as a heuristic strategy to mitigate phylogenetic redundancy, effectively eliminating overrepresented patterns within specific lineages[8]. A higher rJC indicates a greater likelihood that the two KOs co-occur in the same species and may have co-evolved. Additionally, we calculated a hypergeometric p-value (rHyperP) to assess the statistical significance of the rJC between the phylogenetic profiles of each pair of KOs.

Subsequently, for each cooperative KO, a KO was considered to have co-evolved with this cooperative KO if it met three criteria: (i) it must not be another cooperative KO; (ii) the rJC between them must be statistically significant (rHyperP < 0.001); and (iii) the rJC must be larger than 0.25 to exclude KOs that weakly co-occurred with cooperative KOs (Figure S5).

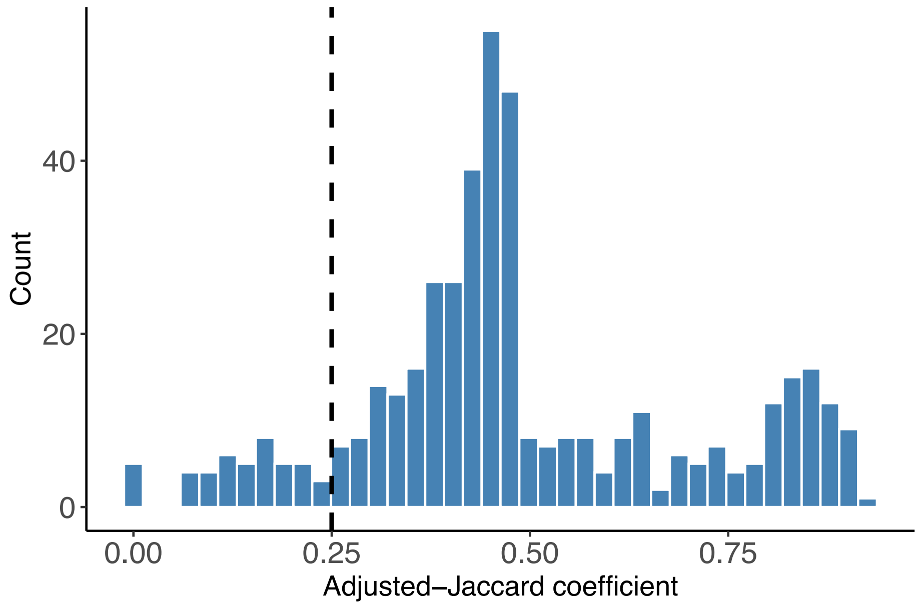


**Figure S5.** Distribution of runs-adjusted Jaccard coefficients (rJCs) between cooperative KOs and candidate co-evolved KOs. The majority of candidate co-evolved KOs and cooperative KOs exhibited rJCs greater than 0.25. We filtered out co-evolved KOs with rJCs smaller than 0.25. This threshold was chosen to exclude

KOs that weakly co-occurred with cooperative KOs, ensuring the analysis included only those with substantial co-evolutionary forces.

Using these criteria, we identified a list of 109 KOs that may have co-evolved with 276 cooperative KOs (Table S14). While co-evolved KOs were defined for each cooperative KO, some cooperative KOs may end up having the same co-evolved KOs (Figure S6).

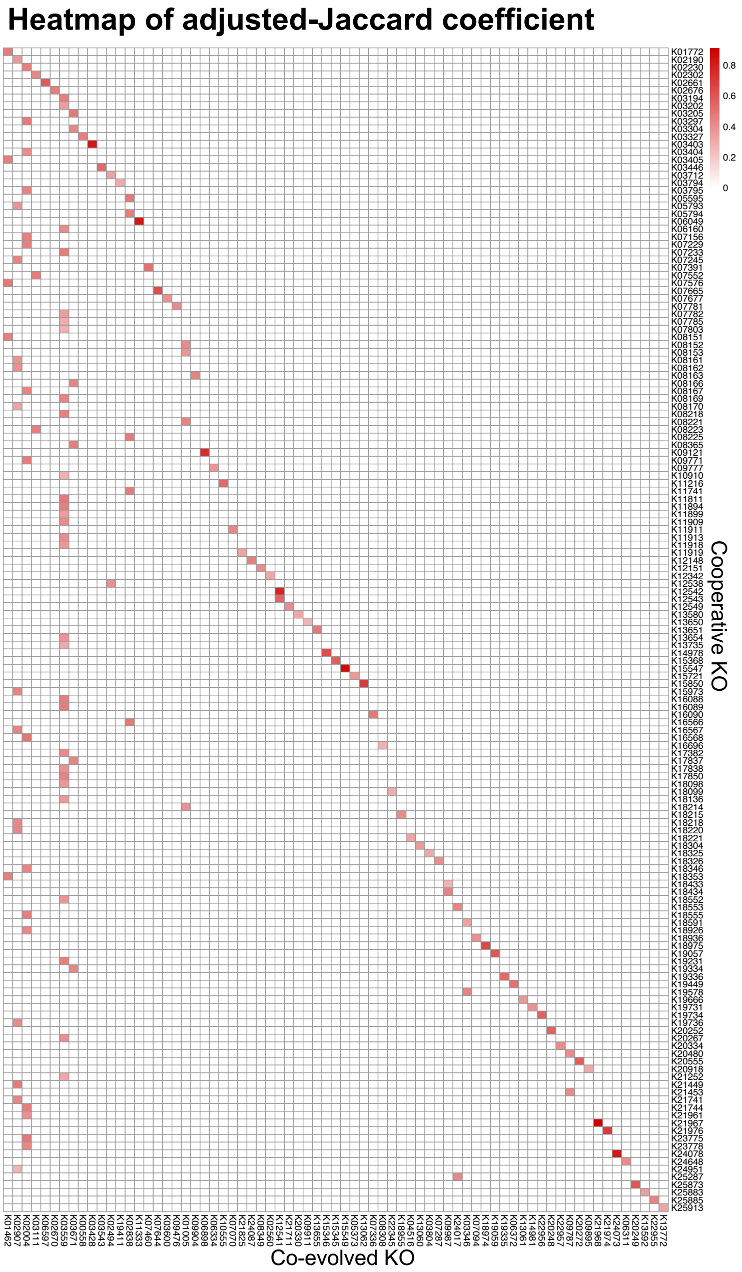

**Figure S6.** This heatmap illustrates the one-to-one runs-adjusted Jaccard coefficients (rJCs) between cooperative KOs (n = 276) and their corresponding co-evolved KOs (n = 109). Some cooperative KOs were associated with the same co-evolved KOs. Certain co-evolved KOs, such as K02004 and K03559, were found to be associated with multiple cooperative KOs. However, in most cases, the coevolution between cooperative KOs and their partners was one-to-one, signifying specific and unique coevolutionary relationships.

b) Examining whether identified co-evolved genes are functionally associated with genes for cooperative traits

If the identified KOs co-evolved with cooperative KOs due to their involvement in similar metabolic pathways, the gain or loss of one gene might directly influence the gain or loss of another gene via functional associations, thereby might affect the role of genes for cooperation in driving niche breadth evolution[2]. We took three steps to test whether cooperative and co-evolved KOs are involved in similar metabolic pathways.

First, we sourced metabolic pathway data for both the cooperative and co-evolved KOs from the KEGG PATHWAY Database[42]. We used the KEGG API (https://rest.kegg.jp/link/ko/pathway) to link KOs (denoted by KO identifiers starting with “ko”) to their corresponding pathways (as represented by pathway identifiers starting with “map”). Vectors containing the metabolic pathways for all cooperative KOs and co-evolved KOs were generated, respectively. Next, we assessed whether cooperative and co-evolved KOs are involved in similar metabolic pathways by computing the Jaccard index between their pathway vectors. Finally, we calculated the P-value of the Jaccard Index using a permutation method, creating a null distribution based on the Jaccard indices between cooperative KOs and 10,000 sets of 109 randomly selected KOs. The p-value was then calculated as the proportion of these permuted Jaccard indices that exceeded the observed Jaccard index between the pathway vectors of cooperative and co-evolved KOs.

Although no significant functional associations were found overall (Figure S7a), we noted that this was primarily because extracellular cooperative KOs and their co-evolved counterparts did not share similar pathways (Figure S7b). In contrast, pathway vectors for intracellular cooperative KOs (20 pathways) and their co-evolved KOs (27 pathways) had a Jaccard Index of 0.237 with a significant permutation p-value of 0.0497, indicating similar pathway involvement (Figure S7c). Therefore, our subsequent analyses focused only on intracellular cooperative genes and their co-evolved partners.

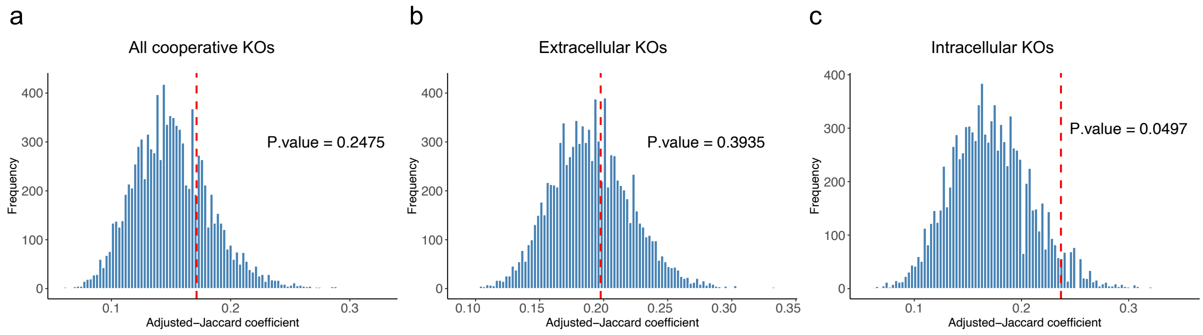


**Figure S7.** Examinations of functional associations between cooperative and co-evolved KOs. The functional associations between cooperative KOs and their co-evolved KOs were estimated using the Jaccard index to measure similarity in metabolic pathways. (a) Overall analysis: The Jaccard index between the metabolic pathways of cooperative KOs and their co-evolved KOs was compared to a null distribution generated from 10,000 sets of randomly selected KOs. Red dashed line represents the observed Jaccard index. The analysis found no significant functional association between cooperative KOs and co-evolved KOs. (b) Analysis of extracellular cooperative KOs: The lack of functional association was primarily due to no significant similarity in metabolic pathways between extracellular cooperative KOs and their co-evolved KOs. (c) Analysis of intracellular cooperative KOs: However, there was a significant similarity in metabolic pathways between intracellular cooperative KOs and their co-evolved partners. This is indicated by a higher Jaccard index compared to the null distribution, with a P-value of 0.0497.

c) Genes that co-evolved with intracellular cooperative genes had limited roles in niche breadth evolution

Next, we explored the role of genes that co-evolved with intracellular cooperative genes in niche breadth evolution. We observed a weak but significant negative correlation between the proportion of co-evolved genes and niche breadth (MCMCglmm; n = 25,785 species; pMCMC < 0.001; Figure S8a; Table S1), contrasting with the expected positive correlation for the number of co-evolved genes (MCMCglmm; n = 25,785 species; pMCMC < 0.001; Figure S8b; Table S1). This finding indicated that although generalists carry more co-evolved genes, their genome sizes increase more rapidly than the number of co-evolved genes.

Additionally, we calculated the possession ratios of cooperative and co-evolved KOs across different niche breadths. For instance, the possession ratio for generalists of a given KO is calculated as the ratio of the number of generalists carrying this KO to the total number of generalists. If a KO is more frequently carried by generalists, the KO is more essential for generalists. We found that the possession ratios of both sets of KOs displayed positive correlations with niche breadth, with higher possession ratios in generalists compared to specialists (Kruskal-Wallis test; cooperative: p-value < 0.001; co-evolved: p-value < 0.001; Figure S8c). However, when checking changes in possession ratios across different niche breadths for each individual KO, we observed that the increase in possession ratios from specialists to generalists was more pronounced for cooperative KOs compared to co-evolved KOs (Figure S8d). Based on the findings from both perspectives, genes co-evolved with intracellular cooperative genes contribute little to niche breadth evolution. The niche breadth evolution was primarily driven by cooperative genes independently.

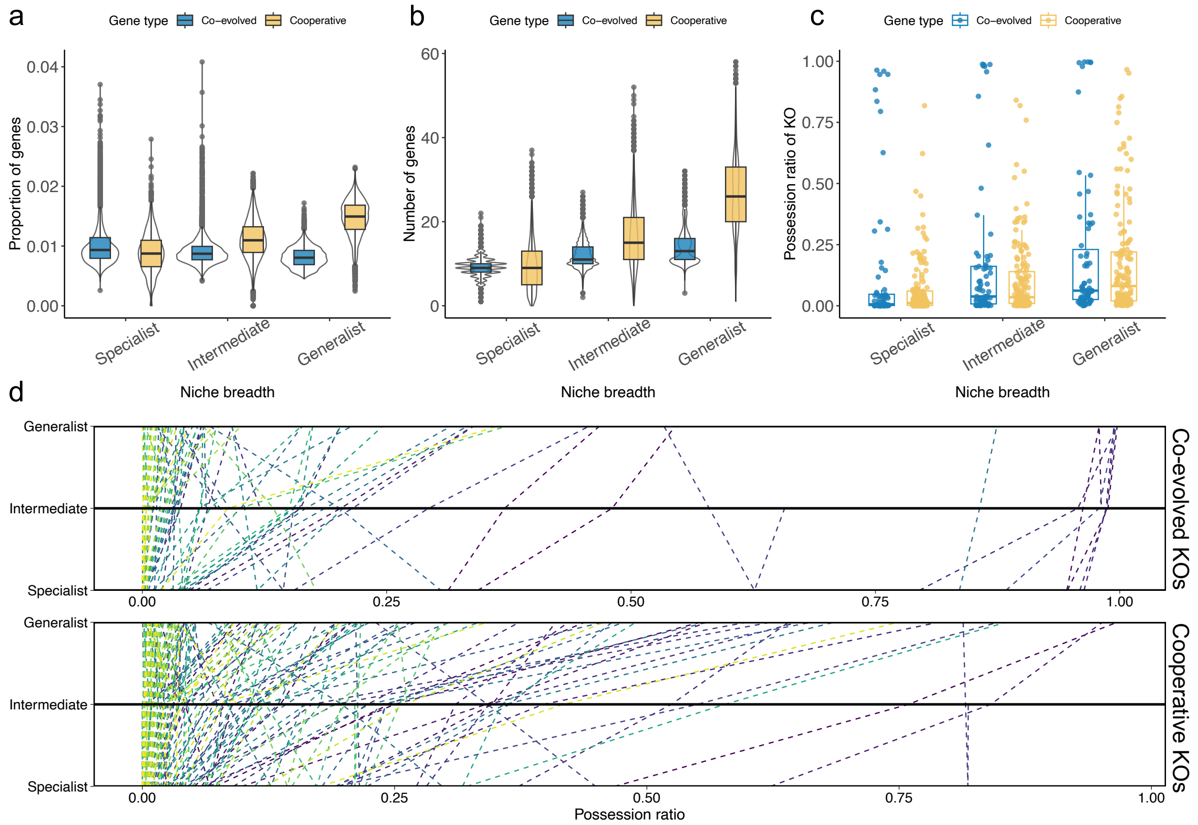


**Figure S8.** Niche breadth evolution was mainly driven by genes for cooperation, not their co-evolved partners. (a) A weak negative correlation between the proportion of co-evolved genes and niche breadth was found, in contrast to intracellular cooperative genes. (b) A positive correlation between the number of co-evolved genes and niche breadth was found, albeit with smaller effect sizes than those observed for intracellular cooperative genes. (c) Positive correlations between the possession ratios of both types of KOs and niche breadth were detected, with generalists showing higher ratios than specialists. For example, possession ratios in generalists are calculated as the number of generalists carrying a specific KO divided by the total number of generalists, applicable across different niche breadths. Each dot represents the possession ratio of a particular KO. (d) Analysis of possession ratio changes across niche breadths for individual KOs revealed more pronounced increases from specialists to generalists in cooperative KOs compared to co-evolved KOs. Dashed lines connect the possession ratios from specialists through intermediates to generalists for each KO.

Overall, our four robustness tests confirmed the finding in the main text that genes for cooperation has a positive and independent role in the niche breadth evolution.

#### Part 2: Robustness tests on the patterns of causal inference

***Test 1:*** *Transition rates analysis confirmed the revealed direction of causality.*

To examine the robustness of the revealed causal relationship between cooperation and niche breadth evolution that a lower proportion of genes for cooperative traits facilitates niche contraction, we conducted an alternative causal inference using transition rates analysis. This method requires the two traits (cooperation and niche breadth) to be binary. Therefore, we binarized these traits (Figure E8) and combined them into a single trait with four states, similar to our approach for fitting MuSSE models in diversification analyses (Figure S9a).

We fitted a Markov (Mk) model for the evolution of this combined trait using the R package *castor*[9,10], and computed the transition rates between states. We focused on two directions of causality: higher levels of cooperation leading to broader niche breadths and lower levels of cooperation leading to narrower niche breadths.

Our analysis found that the transition rate from narrower to broader niche breadths (generalization) was higher in species with more genes for cooperative traits, supporting the causation direction where higher levels of cooperation led to broader niche breadths. Conversely, the transition rate from broader to narrower niche breadths (specialization) was higher in species with fewer genes for cooperative traits, supporting the causation direction where lower levels of cooperation led to narrower niche breadths. Moreover, we found that the rate of specialization was higher than the rate of generalization (Figure S9b). These findings suggested that although both directions of causation can be detected, the direction where lower levels of cooperation led to narrower niche breadth dominated. This was consistent with the results reported in the main text.

We also conducted Bayesian analyses using Markov chain Monte Carlo (MCMC) to examine the posterior distributions of the Mk model parameters using the R package *diversitree* (Figure S9c). The posterior distributions confirmed the results of the one-time Mk model, reinforcing the robustness of our findings.

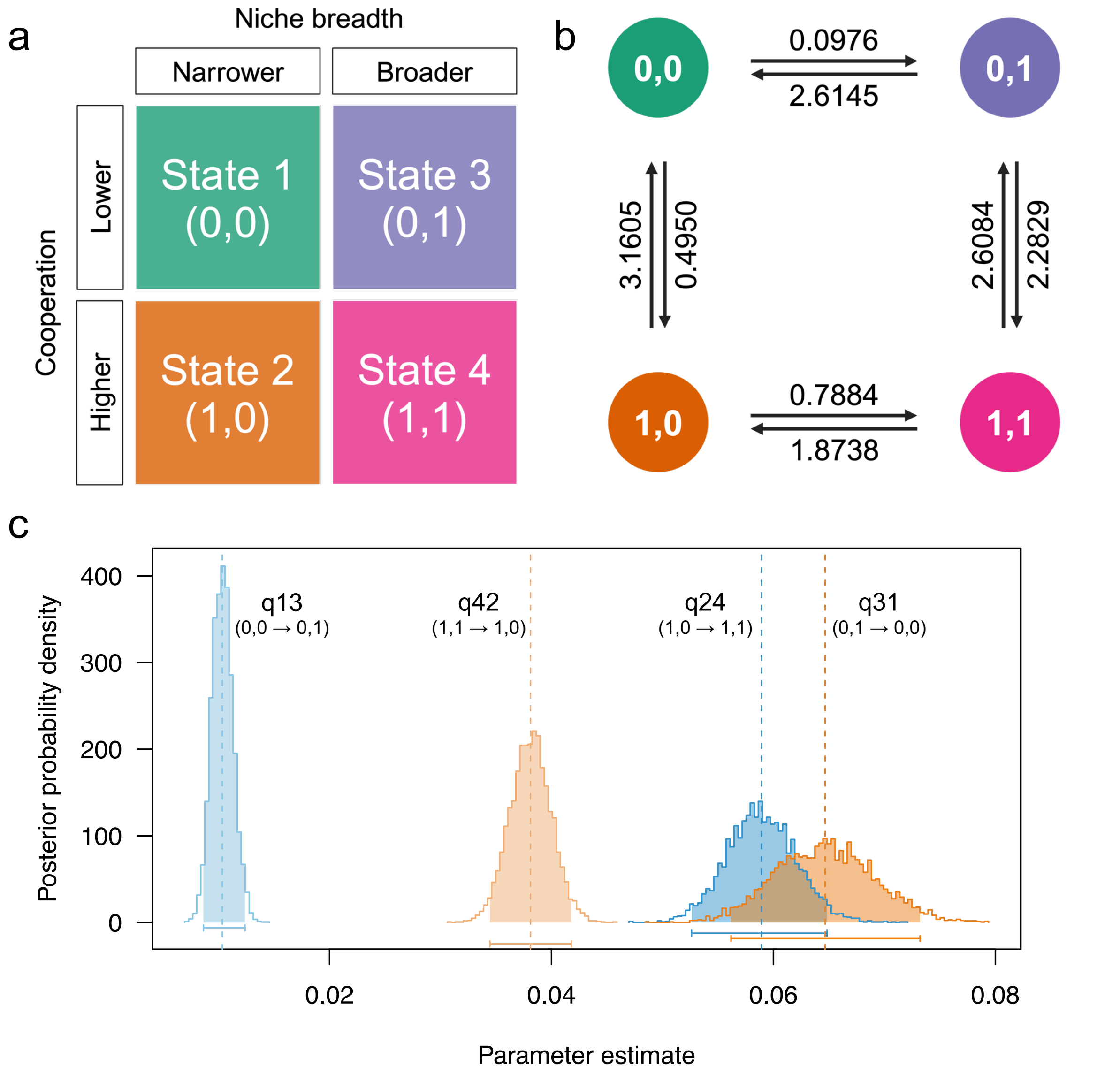

Figure S9. Transition rates analyses as an alternative way of testing causal relationship between cooperation and niche breadth evolution. (a) Like what we did in diversification analyses, we categorized species' niche breadth into "narrower" and "broader" groups, species as having "lower" levels of cooperation or having "higher" levels of cooperation. We then combined the two binary variables into one with four states. (b) We fitted a Markov (Mk) model for the evolution of this combined variable with four states using R package *castor*, and computed the transition rates between states. The numbers alongside the arrows indicate the fitted transition rates. We found that (1) the transition rate from narrower to broader niche breadths (generalization) was higher in species with more cooperative genes ($q_{1,0 \rightarrow 1,1}$) compared to those with fewer cooperative genes ($q_{0,0 \rightarrow 0,1}$); (2) the transition rate from broader to narrower niche breadths (specialization) was higher in species with fewer cooperative genes ($q_{0,1 \rightarrow 0,0}$) compared to those with more cooperative genes ($q_{1,1 \rightarrow 1,0}$); and (3) the rate of specialization ($q_{0,1 \rightarrow 0,0}$) was higher than the rate of generalization ($q_{1,0 \rightarrow 1,1}$). Based on these findings, we concluded that carrying fewer cooperative genes facilitates specialization, which is consistent with previous results analysing the order of evolutionary events. (c) We also conducted Bayesian analyses using Markov chain Monte Carlo (MCMC) to examine the posterior distributions of the Mk model parameters using the R package *diversitree*. We used an exponential prior with a rate of 10, giving a mean of 1/10, and assumed the same prior distribution for all parameters. Using the fitted parameters from the previous Mk model as a starting point, we ran the MCMC for 5,000 iterations, discarding the first 500 samples (10%) to ensure parameter convergence. The posterior probability distributions for the four parameters of interest were presented, with the arithmetic means of the samples indicated by dashed vertical lines, serving as parameter estimates. The bars at the bottom of the distributions and the shaded areas correspond to the 95% credibility intervals. The results were consistent with the previous Mk model findings.

***Test 2:*** *Causal inference using the second optimal threshold for habitat clustering and niche breadth definition.*

In this analysis, we conducted a robustness test on the findings of causal inference using the second optimal threshold for habitat clustering and niche breadth definition (Figure S3). We applied a similar permutation approach as used in the main text to classify species as habitat generalists or specialists based on the number of newly defined habitat clusters they occupied. Species found in one habitat cluster were classified as specialists, those found in eight or more habitat clusters were classified as generalists, and those in between were classified as having “intermediate” niche breadths. Out of the 25,785 species in our dataset, 4,331 were classified as specialists, 14,267 as intermediate species, and 7,187 as generalists (Figure S10, Table S15).

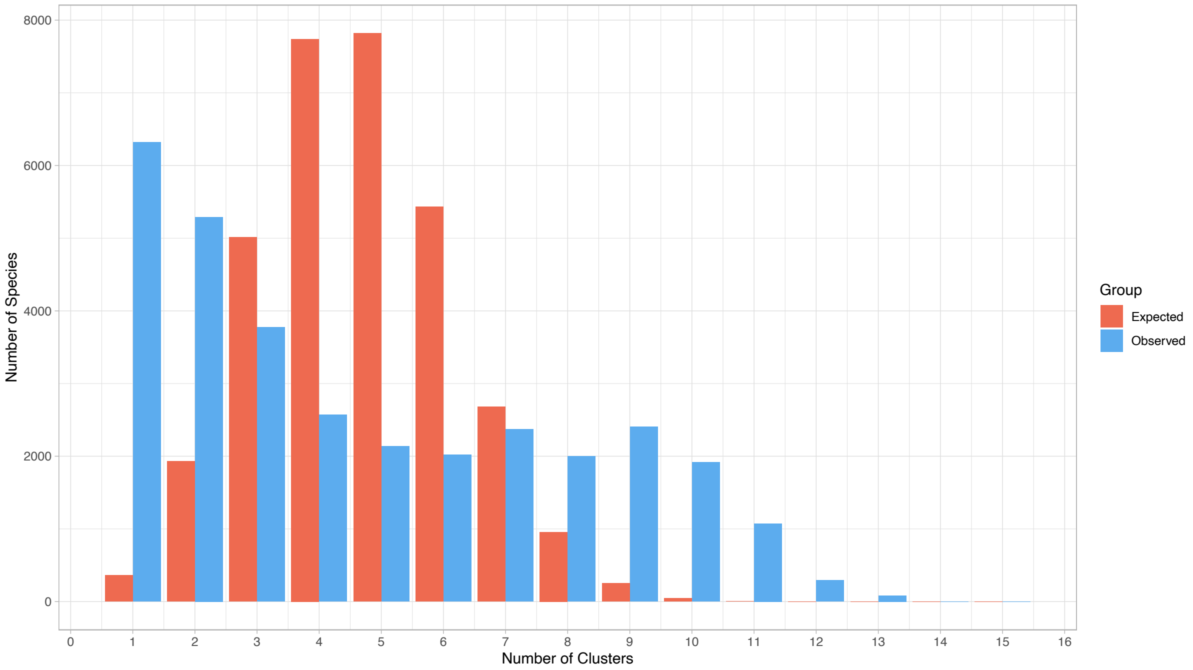


**Figure S10.** Definition of specialists and generalists based on newly defined habitat clusters. Specialists were defined as species found in only one habitat cluster. To define generalists, the observed distribution of niche breadths was compared against an expected distribution generated from 10,000 permutations. Species found in eight or more habitat clusters were identified as generalists, while those found in between were classified as having intermediate niche breadth.

By reconstructing the newly defined niche breadths, we successfully annotated niche breadths for 24,964 ancestral species, with 4,050 specialists, 13,259 species with intermediate niche breadth, and 7,655 generalists (Table S16). Subsequently, we examined the chronological order of cooperation and niche breadth evolution to investigate causality.

First, we investigated whether a higher proportion of genes for cooperation facilitated generalization (Figure 4a). We found that the proportion of genes for cooperation in specialist parents did not significantly affect whether their descendants expanded their niches (MCMCglmm; n = 4,045 specialist ancestral species; pMCMC = 0.694; Figure S11a, Table S1). This suggested that a higher proportion of genes for cooperative traits did not promote niche expansion, consistent with our previous results.

Second, we tested whether a lower proportion of genes for cooperation favoured specialization (Figure 4b). We found that generalist parents whose descendants experienced significant niche contraction had a lower proportion of genes for cooperation (MCMCglmm; n = 7,655 generalist ancestral species; pMCMC < 0.001; Figure S11b, Table S1). This finding supported the direction of causality where a lower proportion of genes for cooperative traits facilitated niche contraction, aligning with our previous finding.

Finally, we simultaneously examined whether generalization promoted an increase in genes for cooperative traits (Figure 4c) or whether specialization facilitated a decrease in such genes (Figure 4d). Specifically, we calculated the change in the proportion of genes for cooperative traits between each ancestral species and the average for its two descendants. Our analyses showed no significant correlation between the ancestral species' niche breadth and the change in the proportion of genes for cooperative traits in their descendants (MCMCglmm; n = 24,963 ancestral species; pMCMC = 0.616; Figure S11c, Table S1). This finding negated causality in both these directions, consistent with our previous finding.

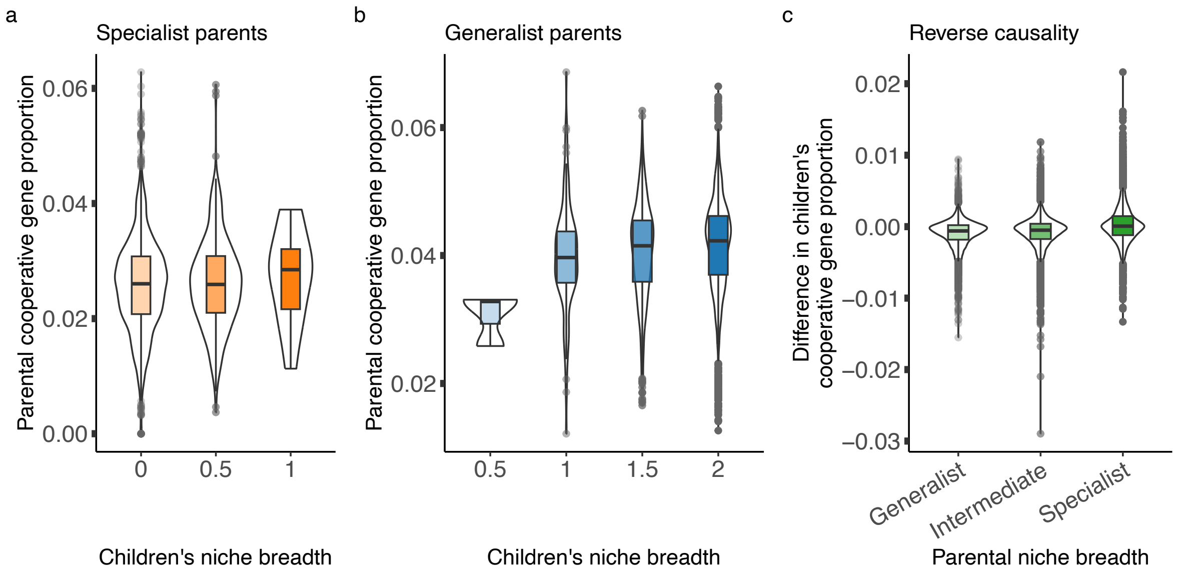


**Figure S11.** Causal inference based on newly defined niche breadth. (a) Generalization: In the evolutionary transition from specialists to generalists, having higher proportions of genes for cooperation did not significantly facilitate generalization. (b) Specialization: Genes for cooperation were found to play a role in the evolutionary transition from generalists to specialists. Specifically, having lower proportions of such genes significantly facilitated specialization. (c) Reverse causality: No significant relationship was observed between the niche breadth of ancestral species and the change in the proportion of genes for cooperation among their descendants. This suggested that neither generalization nor specialization affected the cooperative gene carriage in the offspring, indicating no causal influence in either direction.

Overall, our robustness tests on the causal relationship between cooperation and niche breadth evolution confirmed that the major direction of causation was that a lower proportion of genes for cooperative traits facilitated niche contraction.

#### Part 3: Robustness tests on the patterns of bacterial diversification.

**Test 1:** *The net diversification rates at the family level did not vary significantly across different states.*

We conducted analyses to understand the collective effect of cooperation and niche breadth on the net diversification rates at the family level, defined as speciation rates ($\lambda$) minus extinction rates ($\mu$), in bacteria. The rate parameters were obtained from the MuSSE models used in the main analysis. We observed strong variations in the magnitude of rate parameters across families, with some families exhibiting extremely low net diversification rates ($\lambda - \mu$; Figure S12a). These extreme cases could bias the general pattern when considering all families. To control for this, we introduced two methods of data transformation: (1) $\frac{\lambda-\mu}{\lambda+\mu}$ and (2) $\log(\frac{\lambda}{\mu})$, to reduce the impact of rate magnitude. We found that the net diversification rates at the family level did not vary significantly across different states using either transformation method (MCMCglmm: n = 34 families; Figure S12b, c; Table S1).

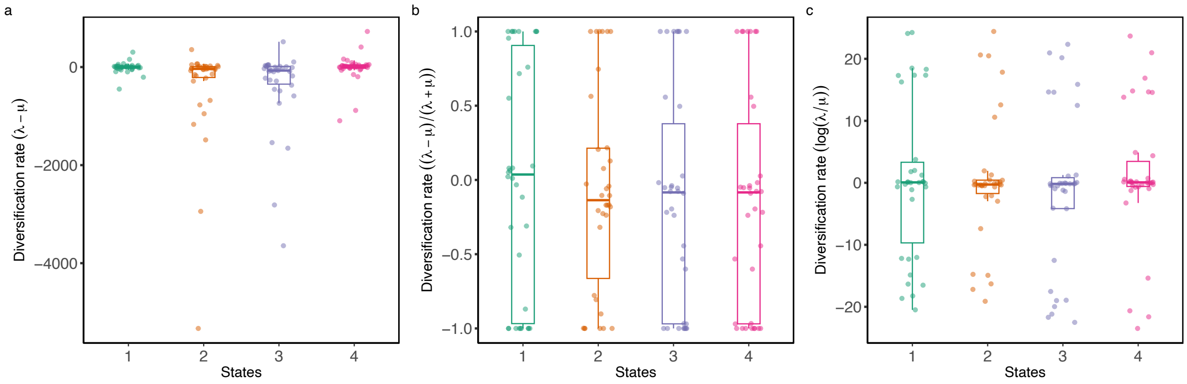


**Figure S12.** Net diversification rates at the family level. (a) Strong variations in the magnitude of rate parameters were detected across families, with some families exhibiting extremely low net diversification rates. To control for these variations, we introduced two methods of data transformation to reduce the magnitude of rate parameters. (b) When diversification rates were calculated as $\frac{\lambda-\mu}{\lambda+\mu}$, we found that the rates at the family level did not vary significantly across different states. (c) When diversification rates were calculated as $\log(\frac{\lambda}{\mu})$, we similarly found that the rates at the family level did not vary significantly across different states.

#### References


1. Arnold, B. J., Huang, I.-T. & Hanage, W. P. Horizontal gene transfer and adaptive evolution in bacteria. *Nat Rev Microbiol* 1–13 (2021) doi:10.1038/s41579-021-00650-4.
2. Tassia, M. G., Whelan, N. V. & Halanych, K. M. Toll-like receptor pathway evolution in deuterostomes. *Proceedings of the National Academy of Sciences* **114**, 7055–7060 (2017).
3. Whelan, F. J., Hall, R. J. & McInerney, J. O. Evidence for Selection in the Abundant Accessory Gene Content of a Prokaryote Pangenome. *Molecular Biology and Evolution* **38**, 3697–3708 (2021).
4. Domingo-Sananes, M. R. & McInerney, J. O. Mechanisms That Shape Microbial Pangenomes. *Trends in Microbiology* **29**, 493–503 (2021).
5. Hall, R. J. *et al. Gene-Gene Relationships in an Escherichia Coli Accessory Genome Are Linked to Function and Mobility*. 2021.03.26.437181 https://www.biorxiv.org/content/10.1101/2021.03.26.437181v1 (2021) doi:10.1101/2021.03.26.437181.
6. Konno, N. & Iwasaki, W. Machine learning enables prediction of metabolic system evolution in bacteria. *Science Advances* **9**, eadc9130 (2023).
7. Tremblay, B. J.-M., Lobb, B. & Doxey, A. C. PhyloCorrelate: inferring bacterial gene–gene functional associations through large-scale phylogenetic profiling. *Bioinformatics* **37**, 17–22 (2021).

8. Cokus, S., Mizutani, S. & Pellegrini, M. An improved method for identifying functionally linked proteins using phylogenetic profiles. *BMC Bioinformatics* **8**, S7 (2007).
9. Louca, S. & Doebeli, M. Efficient comparative phylogenetics on large trees. *Bioinformatics* **34**, 1053–1055 (2018).
10. Louca, S. & Pennell, M. W. A General and Efficient Algorithm for the Likelihood of Diversification and Discrete-Trait Evolutionary Models. *Systematic Biology* **69**, 545–556 (2020).

#### Section 3: Supplementary tables

**Table S1:** MCMCglmm analyses results.

**Table S2:** Results from statistical methods other than MCMCglmm analyses.

**Table S3:** Classification of 114 ProkAtlas habitats into 26 habitat clusters.

**Table S4.** Full list of defined bacterial cooperative KEGG Orthology (KO) terms and their descriptions.

**Table S5.** Raw data for analyses on 25,785 extant species.

**Table S6.** Results of ancestral state reconstruction for niche breadth and gene content.

**Table S7.** Interspecific gene gain and loss rates of 10,060 KOs across 25,785 species.

**Table S8.** Net gain or loss rates for 579 cooperative KOs.

**Table S9.** Intraspecific gene gain and loss rates of cooperative genes in 171 species.

**Table S10.** List of families on which diversification analysis was based.

**Table S11.** Annotations of ProkAtlas habitat preferences for 32,262 species.

**Table S12.** Annotations of habitat cluster presence/absence for 32,262 species.

**Table S13.** Silhouette indices at different sequence identify thresholds and number of clusters when clustering method = "ward.D2".

**Table S14.** Metabolic pathway information and runs-adjusted Jaccard coefficient (rJC) between cooperative KOs and corresponding co-evolved KOs.

**Table S15.** Annotations of alternatively defined habitat clusters for 32,262 species.

**Table S16.** Results of ancestral state reconstruction for newly defined niche breadth.
