## Supplementary material for "Cooperation shapes bacterial niche breadth evolution and patterns of diversification": Supplmentary tables: TableS1.docx

**Table S1**. MCMCglmm analyses results.

| **Model description** | **Term** | **Posterior mean** | **l-95% CI** | **u-95% CI** | **pMCMC** | **Sig.** | **Sample size** |
| --- | --- | --- | --- | --- | --- | --- | --- |
| Extant species: Proportion of cooperative genes ~ Niche breadth (overall) | Niche Breadth | 0.0004236 | 0.0003848 | 0.0004659 | < 0.001 | *** | 25,785 species |
| Extant species: Proportion of cooperative genes ~ Niche breadth (extracellular protein) | Niche Breadth | 0.0001273 | 0.0001062 | 0.0001475 | < 0.001 | *** | 25,785 species |
| Extant species: Proportion of cooperative genes ~ Niche breadth (antibiotic degradation) | Niche Breadth | 0.0001435 | 0.0001300 | 0.0001580 | < 0.001 | *** | 25,785 species |
| Extant species: Proportion of cooperative genes ~ Niche breadth (biofilm formation) | Niche Breadth | 4.442e-05 | 3.219e-05 | 5.573e-05 | < 0.001 | *** | 25,785 species |
| Extant species: Proportion of cooperative genes ~ Niche breadth (quorum sensing) | Niche Breadth | 8.352e-06 | 3.893e-06 | 1.283e-05 | < 0.001 | *** | 25,785 species |
| Extant species: Proportion of cooperative genes ~ Niche breadth (secretion system) | Niche Breadth | 8.707e-05 | 6.512e-05 | 1.085e-04 | < 0.001 | *** | 25,785 species |
| Extant species: Proportion of cooperative genes ~ Niche breadth (siderophores) | Niche Breadth | 5.224e-05 | 4.031e-05 | 6.329e-05 | < 0.001 | *** | 25,785 species |
| *Causality: Whether having a higher proportion of cooperative genes facilitated generalization.* Ancestral species: Children’s niche breadth ~ Ancestral proportion of cooperative genes | Proportion of cooperative genes | -3.677 | -20.075 | 10.671 | 0.637 | ns | 3,832 species |
| *Causality: Whether having a lower proportion of cooperative genes facilitated specialization.*  Ancestral species: Ancestral proportion of cooperative genes ~ Children’s niche breadth | 1 vs. 0.5 | 0.005147 | 0.002644 | 0.007909 | < 0.001 | *** | 9,173 species |
|  | 1 vs. 1.5 | -0.0001393 | -0.000741 | 0.0004967 | 0.6630 | ns |  |
|  | 2 vs. 1.5 | 0.0004928 | 0.0003338 | 0.0006478 | < 0.001 | *** |  |
| *Causality: Whether generalization promoted an increase in cooperative genes or whether specialization facilitated a decrease in such genes.*  Ancestral species: Children’s proportion of cooperative genes ~ Ancestral niche breadth | Generalists  vs.  Intermediate | 3.196e-05 | -2.994e-05 | 1.004e-04 | 0.3323 | ns | 24,590 species |
|  | Generalists  vs.  Specialists | 2.363e-05 | -7.803e-05 | 1.284e-04 | 0.6434 | ns |  |
| Overall: propensity of cooperative genes being accessory genes in pangenome ($\pi$) ~ 1 | Intercept | 0.04383 | 0.01823 | 0.06955 | < 0.001 | *** | 171 species |
| Biofilm formation: propensity of cooperative genes being accessory genes in pangenome ($\pi$) ~ 1 | Intercept | 0.01675 | -0.03178 | 0.06241 | 0.478 | ns | 171 species |
| Quorum sensing: propensity of cooperative genes being accessory genes in pangenome ($\pi$) ~ 1 | Intercept | -0.007003 | -0.071702 | 0.059611 | 0.82 | ns | 171 species |
| Siderophores: propensity of cooperative genes being accessory genes in pangenome ($\pi$) ~ 1 | Intercept | 0.006735 | -0.027307 | 0.043184 | 0.677 | ns | 171 species |
| Secretion system: propensity of cooperative genes being accessory genes in pangenome ($\pi$) ~ 1 | Intercept | 0.15858 | 0.01523 | 0.29784 | 0.037 | * | 171 species |
| Extracellular protein: propensity of cooperative genes being accessory genes in pangenome ($\pi$) ~ 1 | Intercept | 0.016907 | -0.005371 | 0.037312 | 0.129 | ns | 171 species |
| Antibiotic degradation: propensity of cooperative genes being accessory genes in pangenome ($\pi$) ~ 1 | Intercept | 0.04245 | 0.01213 | 0.07232 | 0.0132 | * | 171 species |
| Diversification: Speciation rate (lambda) ~ Level of cooperation | lambda1  vs.  lambda0 | 1.4385 | -1.2756 | 4.2161 | 0.304 | ns | 46 families |
| Diversification: Extinction rate (mu) ~ Level of cooperation | mu1  vs.  mu0 | 4.5208 | 1.7254 | 7.2575 | 0.0034 | ** | 46 families |
| Diversification: Speciation rate (lambda) ~ Niche breadth | lambda1  vs.  lambda0 | 3.70496 | 1.04497 | 6.30693 | 0.00383 | ** | 45 families |
| Diversification: Extinction rate (mu) ~ Niche breadth | mu1  vs.  mu0 | -0.2047 | -1.9081 | 1.5400 | 0.826 | ns | 45 families |
| Diversification: Speciation rate (lambda) ~ Level of cooperation* Niche breadth (4 states) | lambda2  vs.  lambda1 | 4.0081 | 0.3673 | 7.6558 | 0.0319 | * | 34  families |
|  | lambda3  vs.  lambda1 | 2.3051 | -1.5867 | 5.7906 | 0.2213 | ns |  |
|  | lambda4  vs.  lambda2 | 0.03161 | -3.71597 | 3.48762 | 0.9817 | ns |  |
|  | lambda4  vs.  lambda3 | 1.6628 | -1.8508 | 5.2583 | 0.357 | ns |  |
| Diversification: Extinction rate (mu) ~ Level of cooperation* Niche breadth (4 states) | mu2  vs.  mu1 | 4.23744 | 0.25501 | 8.31063 | 0.0383 | * | 34  families |
|  | mu3  vs.  mu1 | 4.11332 | -0.01615 | 8.10259 | 0.0468 | * |  |
|  | mu4  vs.  mu2 | -0.96054 | -5.11901 | 2.97912 | 0.6357 | ns |  |
|  | mu4  vs.  mu3 | -0.88684 | -4.96849 | 3.01269 | 0.6438 | ns |  |
| **Supplementary results** Extant species: Proportion of cooperative genes ~ Newly defined niche breadth (overall) | Niche Breadth | 0.0004540 | 0.0004117 | 0.0005021 | < 0.001 | *** | 25,785 species |
| **Supplementary results**  Extant species: Proportion of cooperative genes ~ Newly defined niche breadth (extracellular protein) | Niche Breadth | 0.0001352 | 0.0001115 | 0.0001575 | < 0.001 | *** | 25,785 species |
| **Supplementary results**  Extant species: Proportion of cooperative genes ~ Newly defined niche breadth (antibiotic degradation) | Niche Breadth | 0.0001494 | 0.0001349 | 0.0001657 | < 0.001 | *** | 25,785 species |
| **Supplementary results**  Extant species: Proportion of cooperative genes ~ Newly defined niche breadth (biofilm formation) | Niche Breadth | 4.573e-05 | 3.213e-05 | 5.812e-05 | < 0.001 | *** | 25,785 species |
| **Supplementary results**  Extant species: Proportion of cooperative genes ~ Newly defined niche breadth (quorum sensing) | Niche Breadth | 8.468e-06 | 3.564e-06 | 1.324e-05 | < 0.001 | *** | 25,785 species |
| **Supplementary results**  Extant species: Proportion of cooperative genes ~ Newly defined niche breadth (secretion system) | Niche Breadth | 9.526e-05 | 7.233e-05 | 1.206e-04 | < 0.001 | *** | 25,785 species |
| **Supplementary results**  Extant species: Proportion of cooperative genes ~ Newly defined niche breadth (siderophores) | Niche Breadth | 6.191e-05 | 4.962e-05 | 7.454e-05 | < 0.001 | *** | 25,785 species |
| **Supplementary results** Extant species: Number of cooperative genes ~ Niche breadth (overall) | Niche Breadth | 1.350 | 1.257 | 1.435 | < 0.001 | *** | 25,785 species |
| **Supplementary results**  Extant species: Number of cooperative genes ~ Niche breadth (extracellular protein) | Niche Breadth | 0.5120 | 0.4749 | 0.5506 | < 0.001 | *** | 25,785 species |
| **Supplementary results**  Extant species: Number of cooperative genes ~ Niche breadth (antibiotic degradation) | Niche Breadth | 0.3300 | 0.3067 | 0.3529 | < 0.001 | *** | 25,785 species |
| **Supplementary results**  Extant species: Number of cooperative genes ~ Niche breadth (biofilm formation) | Niche Breadth | 0.1508 | 0.1338 | 0.1680 | < 0.001 | *** | 25,785 species |
| **Supplementary results**  Extant species: Number of cooperative genes ~ Niche breadth (quorum sensing) | Niche Breadth | 0.01532 | 0.01087 | 0.01969 | < 0.001 | *** | 25,785 species |
| **Supplementary results**  Extant species: Number of cooperative genes ~ Niche breadth (secretion system) | Niche Breadth | 0.2299 | 0.1910 | 0.2667 | < 0.001 | *** | 25,785 species |
| **Supplementary results**  Extant species: Number of cooperative genes ~ Niche breadth (siderophores) | Niche Breadth | 0.1501 | 0.1331 | 0.1679 | < 0.001 | *** | 25,785 species |
| **Supplementary results** Extant species: Niche breadth with 3 categories ~ Proportion of cooperative genes (overall) | Proportion of cooperative genes | -37.272 | -40.656 | -33.508 | < 0.001 | *** | 25,785 species |
| **Supplementary results**  Extant species: Niche breadth with 3 categories ~ Proportion of cooperative genes (extracellular protein) | Proportion of cooperative genes | -39.633 | -48.716 | -31.146 | < 0.001 | *** | 25,785 species |
| **Supplementary results**  Extant species: Niche breadth with 3 categories ~ Proportion of cooperative genes (antibiotic degradation) | Proportion of cooperative genes | -124.276 | -136.560 | -113.207 | < 0.001 | *** | 25,785 species |
| **Supplementary results**  Extant species: Niche breadth with 3 categories ~ Proportion of cooperative genes (biofilm formation) | Proportion of cooperative genes | -38.199 | -50.374 | -24.981 | < 0.001 | *** | 25,785 species |
| **Supplementary results**  Extant species: Niche breadth with 3 categories ~ Proportion of cooperative genes (quorum sensing) | Proportion of cooperative genes | -155.280 | -210.763 | -101.999 | < 0.001 | *** | 25,785 species |
| **Supplementary results**  Extant species: Niche breadth with 3 categories ~ Proportion of cooperative genes (secretion system) | Proportion of cooperative genes | -23.303 | -29.141 | -17.454 | < 0.001 | *** | 25,785 species |
| **Supplementary results**  Extant species: Niche breadth with 3 categories ~ Proportion of cooperative genes (siderophores) | Proportion of cooperative genes | -74.578 | -86.105 | -63.526 | < 0.001 | *** | 25,785 species |
| **Supplementary results** *Causality: Whether having a higher proportion of cooperative genes facilitated generalization.* Ancestral species: Children’s niche breadth (newly defined) ~ Ancestral proportion of cooperative genes | Proportion of cooperative genes | -3.505 | -23.691 | 17.772 | 0.694 | ns | 4045  species |
| **Supplementary results** *Causality: Whether having a lower proportion of cooperative genes facilitated specialization.*  Ancestral species: Children’s niche breadth (newly defined) ~ Ancestral proportion of cooperative genes | Proportion of cooperative genes | 30.2636 | 19.4881 | 38.9997 | < 0.001 | *** | 7655 species |
| **Supplementary results** *Causality: Whether generalization promoted an increase in cooperative genes or whether specialization facilitated a decrease in such genes.*  Ancestral species: Ancestral niche breadth ~ Children’s proportion of cooperative genes | Proportion of cooperative genes | 1.640 | -4.776 | 7.659 | 0.616 | ns | 24,963 species |
| **Supplementary results** Diversification: $\frac{\lambda-\mu}{\lambda+\mu}$~ Level of cooperation* Niche breadth (4 states) | div2  vs.  div1 | -0.076616 | -0.404487 | 0.268571 | 0.647 | ns | 34  families |
|  | div3  vs.  div1 | -0.135415 | -0.450446 | 0.209630 | 0.440 | ns | 34  families |
|  | div4  vs.  div2 | -0.05644 | -0.37678 | 0.27480 | 0.743 | ns | 34  families |
|  | div4  vs.  div3 | 0.000527 | -0.304127 | 0.346427 | 0.993 | ns | 34  families |
| **Supplementary results** Diversification: $\log(\frac{\lambda}{\mu})$ ~ Level of cooperation* Niche breadth (4 states) | div2  vs.  div1 | -0.1582 | -5.3733 | 5.4112 | 0.963 | ns | 34  families |
|  | div3  vs.  div1 | -1.8018 | -7.2811 | 3.7426 | 0.529 | ns | 34  families |
|  | div4  vs.  div2 | 1.0024 | -4.1430 | 6.7783 | 0.731 | ns | 34  families |
|  | div4  vs.  div3 | 2.543 | -2.687 | 8.401 | 0.372 | ns | 34  families |
